# Transcription factor Sp1 regulates mitotic fidelity through Aurora B kinase-mediated condensin I localization

**DOI:** 10.1101/2020.06.19.158030

**Authors:** Samuel Flashner, Michelle Swift, Aislinn Sowash, Jane Azizkhan-Clifford

## Abstract

Mitotic chromosome assembly is essential for faithful chromosome segregation. Despite their salient role directing interphase chromatin organization, little is known about how transcription factors mediate this process during mitosis. Here, we characterize a mitosis-specific role for transcription factor specificity protein 1 (Sp1). Sp1 localizes to mitotic centromeres and auxin-induced rapid Sp1 degradation results in chromosome segregation errors and aberrant mitotic progression. These defects are driven by anomalous mitotic chromosome assembly. Sp1 degradation results in chromosome condensation defects through reduced condensin complex I localization. Sp1 also mediates the localization and activation of Aurora B kinase early in mitosis, which is essential for condensin complex I recruitment. Underscoring the clinical significance of our findings, aberrant Sp1 expression correlates with aneuploidy in several human cancers, including kidney renal papillary cell carcinoma, ovarian serous cystadenocarcinoma, mesothelioma, cholangiocarcinoma, and hepatocellular carcinoma. Our results suggest that Sp1 protects genomic integrity during mitosis by promoting chromosome assembly.

## Introduction

Chromosome segregation errors are detectable in up to 90% of solid tumors (Weaver & Cleveland, 2006). Persistent chromosome missegregation is associated with poor patient prognosis, decreased patient survival, intrinsic multidrug resistance, and increased intratumoral heterogeneity (Bakhoum & Landau, 2017; A. J. X. Lee et al., 2011; McGranahan, Burrell, Endesfelder, Novelli, & Swanton, 2012). Characterizing the molecular mechanisms governing chromosome segregation is therefore a clinical imperative. Mitotic chromosome assembly is essential for mitotic fidelity, yet the factors mediating this process are incompletely characterized. Transcription factors are the second most abundant class of gene in humans and mediate interphase chromatin dynamics (Venter et al., 2001). Despite the preponderance of circumstantial evidence linking transcription factors to mitotic chromosome assembly, their role in this process is undefined.

Dysregulation of transcription factors is associated with increased chromosome segregation errors in a variety of contexts (Astrinidis et al., 2010; Ishak, Coschi, Roes, & Dick, 2017; H.-S. Lee et al., 2018; Rohrberg et al., 2020; Weiler et al., 2017). However, several barriers have obfuscated the direct role of transcription factors in chromosome segregation. Until recently, transcription factors were believed to be globally evicted from mitotic chromatin and therefore were overlooked during cell division (Ginno, Burger, Seebacher, Iesmantavicius, & Schübeler, 2018; Martinez-Balbbs, Dey, Rabindran, Ozato, & Wu, 1995; Teves et al., 2016). Mitotic retention of transcription factors has been implicated in mitotic bookmarking which is required for the maintenance of transcriptional programs into G1 (Caravaca et al., 2013; Kadauke et al., 2012; Teves et al., 2016). However, how retention of these factors contributes to mitotic fidelity has not been studied. Additionally, standardly used loss-of-function molecular techniques slowly deplete protein levels over several cell divisions, thereby interrupting their transcriptional programs and confounding their mitosis-specific effects. Faithful characterization of mitotic transcription factor function therefore requires more rapid protein depletion immediately prior to mitosis. Recent advancements in degron technology have enabled progress in this field; however, few studies have examined the role of individual transcription factors during mitosis (Natsume & Kanemaki, 2017). These studies require an abundance of resources; therefore, which transcription factors are evaluated needs to be judiciously selected. The ideal candidates have been implicated in chromosome segregation through poorly defined mechanisms and localize to mitotic chromatin for poorly defined purpose. Specificity protein 1 (Sp1) is a ubiquitously expressed mammalian transcription factor that protects genomic integrity through diverse mechanisms (Astrinidis et al., 2010; Beishline et al., 2012; Torabi et al., 2018). RNAi-mediated depletion of Sp1 results in chromosome missegregation (Astrinidis et al., 2010). While recent evidence suggests that Sp1 localizes to mitotic chromatin, its precise role during mitosis is undefined (Ginno et al., 2018; Teves et al., 2016). Sp1 is therefore an ideal candidate for the current study. Here, we interrogate the mitosis-specific role of the transcription factor Sp1 by rapidly degrading it immediately prior to mitotic entry.

Transcription factors direct the organization and assembly of interphase chromatin; however, their role in these processes during mitosis is undefined (Seungsoo Kim & Shendure, 2019). Errorless chromosome segregation requires the proper assembly of mitotic chromosomes by condensin complexes I and II (Hirano, 2016). Both condensin complexes localize along the chromosome arms and reorganize chromatin early in mitosis. Condensin complexes I and II have distinct functions and dynamics during mitosis (Hirota, Gerlich, Koch, Ellenberg, & Peters, 2004). Condensin complex II stably associates with chromosomes during DNA replication and axially compacts chromatin in prophase (Ono, Yamashita, & Hirano, 2013; Walther et al., 2018). Condensin complex I dynamically associates with mitotic chromosome arms after nuclear envelope breakdown and drives lateral chromosome compaction during mitosis (Gerlich, Hirota, Koch, Peters, & Ellenberg, 2006; Walther et al., 2018). While both condensin complexes contain the chromosomal ATPases SMC2 and SMC4, the complexes are composed of unique subunits: nCAP-D2, nCAP-G, and nCAP-H comprise condensin I while nCAP-D3, nCAP-G2, and nCAP-H2 comprise condensin II (Hirano, 2012). Preventing the association of either condensin complex I or II results in chromosome segregation errors in tissue culture cells (Booth et al., 2016; Samoshkin et al., 2009). Further, condensin complex function is clinically relevant; defective chromosome condensation is associated with leukemia and other disease states (Molina et al., 2020; Woodward et al., 2016). Despite the importance of condensin-mediated mitotic chromatin assembly, little is known about the molecular mechanisms regulating their stability and localization to mitotic chromosomes.

Aurora B kinase is required for condensin-mediated mitotic chromosome assembly through poorly defined mechanisms (Lipp, Hirota, Poser, & Peters, 2007; Takemoto et al., 2007). Aurora B forms the chromosomal passenger complex (CPC) with Borealin, INCENP, and Survivin, which is required for chromosome condensation, correcting microtubule/kinetochore attachments, and cytokinesis (Carmena, Wheelock, Funabiki, & Earnshaw, 2012). CPC activity and function are tightly coupled to its spatiotemporal localization. Early in mitosis, the CPC localizes to chromosome arms in a process required for chromosome condensation. However, little is known about how the CPC is targeted to the chromatin in prophase. Loss of aurora B early in mitosis results in a variety of deleterious phenotypes that phenocopy chromosome condensation defects (Molina et al., 2020; Samoshkin et al., 2009). More work is needed to fully characterize the aurora B – condensin signaling axis in chromosome segregation.

In this study, we find that transcription factor Sp1 mediates Aurora B localization to mitotic arms and the recruitment of condensin complex I. Auxin-induced rapid Sp1 degradation immediately prior to mitosis results in mitotic defects and chromosome segregation errors. Corroborating these results, we find that aberrant Sp1 expression correlates with aneuploidy in human cancers. Taken together, we implicate a ubiquitously expressed transcription factor in a clinically relevant, yet poorly understood phenomenon. These data challenge the current paradigm that transcription factors have no direct role in promoting mitotic fidelity.

## Results

### Sp1 dynamically localizes to mitotic centromeres

Sp1 was recently shown to localize to mitotic chromatin (Ginno et al., 2018; Teves et al., 2016). While Sp1 is predicted to function as a bookmarker at mitotic chromatin, this hypothesis has not been tested (Ginno et al., 2018; Teves et al., 2016). In order to evaluate this possibility, we first examined Sp1 localization dynamics during mitosis. While previous work demonstrated that Sp1 localizes along entire metaphase chromosome arms, we found that Sp1 colocalizes with metaphase centromeres dynamically during mitosis (Fig. 1A-D) (Teves et al., 2016). Sp1 colocalizes with CENP-A, a marker of the centromere, in metaphase retinal pigment epithelial (RPE-1) cells (Fig. 1A-D). Sp1’s peak positional intensity is slightly offset from CENP-A’s, indicating that Sp1 localizes to the centromeres and may be localizing to the adjacent pericentromeres (Fig. 1C). We next examined Sp1 localization at different stages of mitosis in normal human diploid fibroblasts (NHDFs). Sp1 is detectable at centromeres during prophase (Fig. 1D). While Sp1 is detectable at nearly every centromere during metaphase (Fig. 1C), by anaphase Sp1 does not appear to overlap with CENP-A (Fig. 1D. During interphase, there is no apparent localization of Sp1 to the centromeric region, indicating that this dynamic localization occurs specifically in mitotic cells. Together, these data indicate that Sp1 dynamically localizes to the centromere during mitotic progression. Because Sp1 is evicted from the centromere prior to entry into G1, we predict that Sp1 is not functioning as a mitotic bookmarker at this region.

**Fig. 1.**
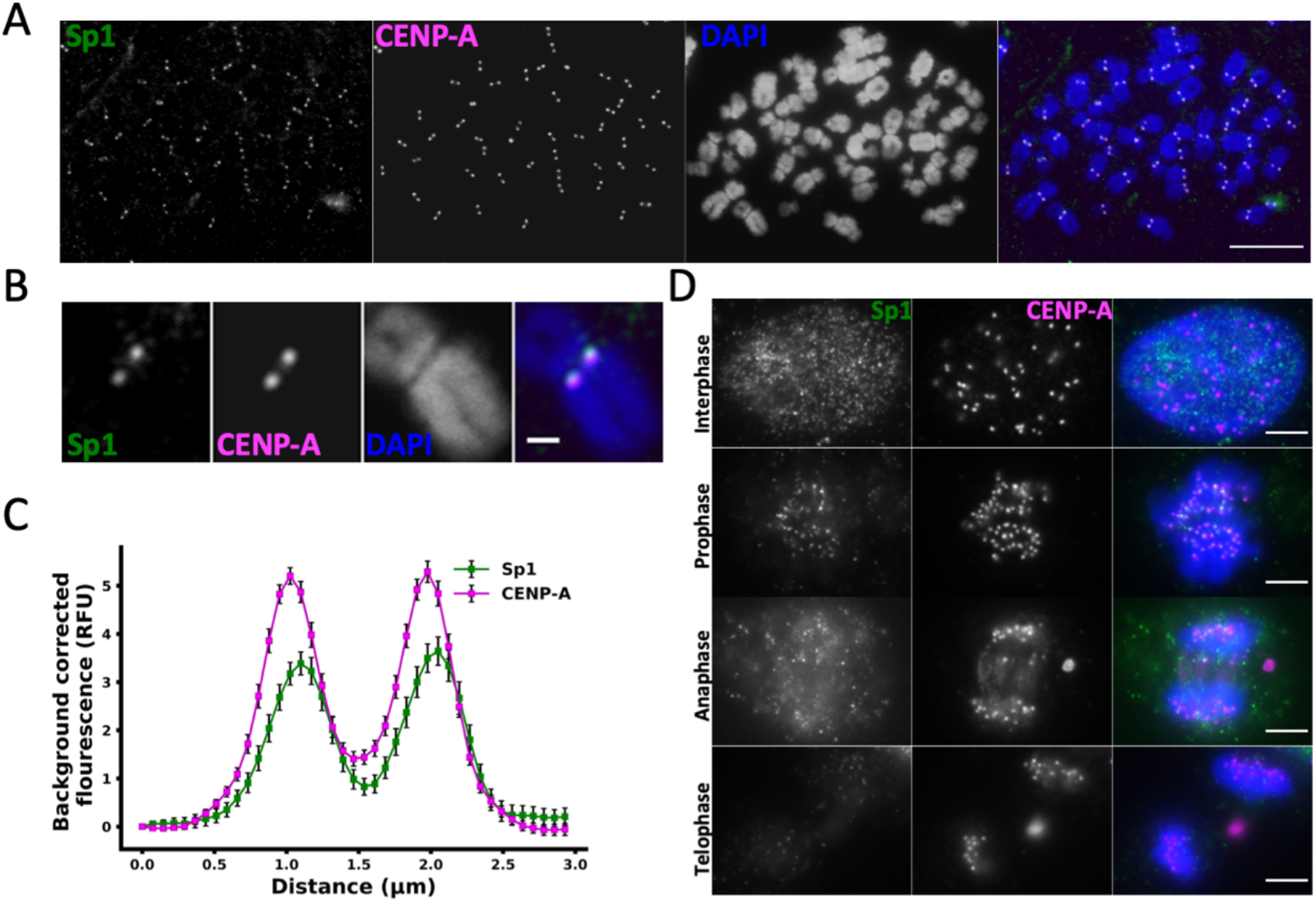
Sp1 localizes to mitotic centromeres. **(A)** RPE-1 cells were arrested in metaphase, spread onto a glass slide, and stained for Sp1 and CENP-A. Scale bar = 10 μm. White box indicates the inset for **(B).** Scale bar = 1 μm. **(C)** Plot profile of the positional intensity of Sp1 and CENP-A. Each point represents the average positional intensity in 50 centromere pairs. Error bars represent the standard error of the mean (s.e.m.). **(D)** NHDFs were fixed and stained for Sp1 and CENP-A. Their cell cycle phase was scored based on the conformation of their chromosomes. Scale bar = 5 μm.

### Sp1 regulates chromosome segregation during mitosis

We next wanted to characterize the mitosis-specific function for Sp1. Because siRNA-mediated Sp1 depletion results in chromosome segregation errors, we hypothesized that Sp1 functions during mitosis to regulate chromosome segregation. However, characterizing the mitotic function of transcription factors is challenging; RNAi or CRISPR-mediated depletion of transcription factors is too slow to separate their mitosis-specific functions from their interphase transcriptional programs. We overcame this limitation by utilizing an auxin-inducible degron (AID) to rapidly deplete the ubiquitously expressed transcription factor Sp1 in the chromosomally-stable cell line RPE-1 (mAID-Sp1) (Fig. 2A) (Clift et al., 2017; Holland, Fachinetti, Han, & Cleveland, 2012; Nishimura, Fukagawa, Takisawa, Kakimoto, & Kanemaki, 2009). We were able to reduce Sp1 protein levels by treating these cells with auxin for one hour (Fig. 2B). In order to determine if we could degrade Sp1 on mitotic chromosomes, we treated mAID-Sp1 cells with auxin for two hours and then arrested them in metaphase with colcemid for an additional two hours. Following this treatment, Sp1 was depleted at mitotic centromeres (Fig. 2C, D). This treatment strategy ensures that all visualized AID-Sp1 cells enter mitosis shortly after Sp1 has been degraded (Fig. 2C), isolating the mitosis-specific effects of Sp1.

**Fig. 2.**
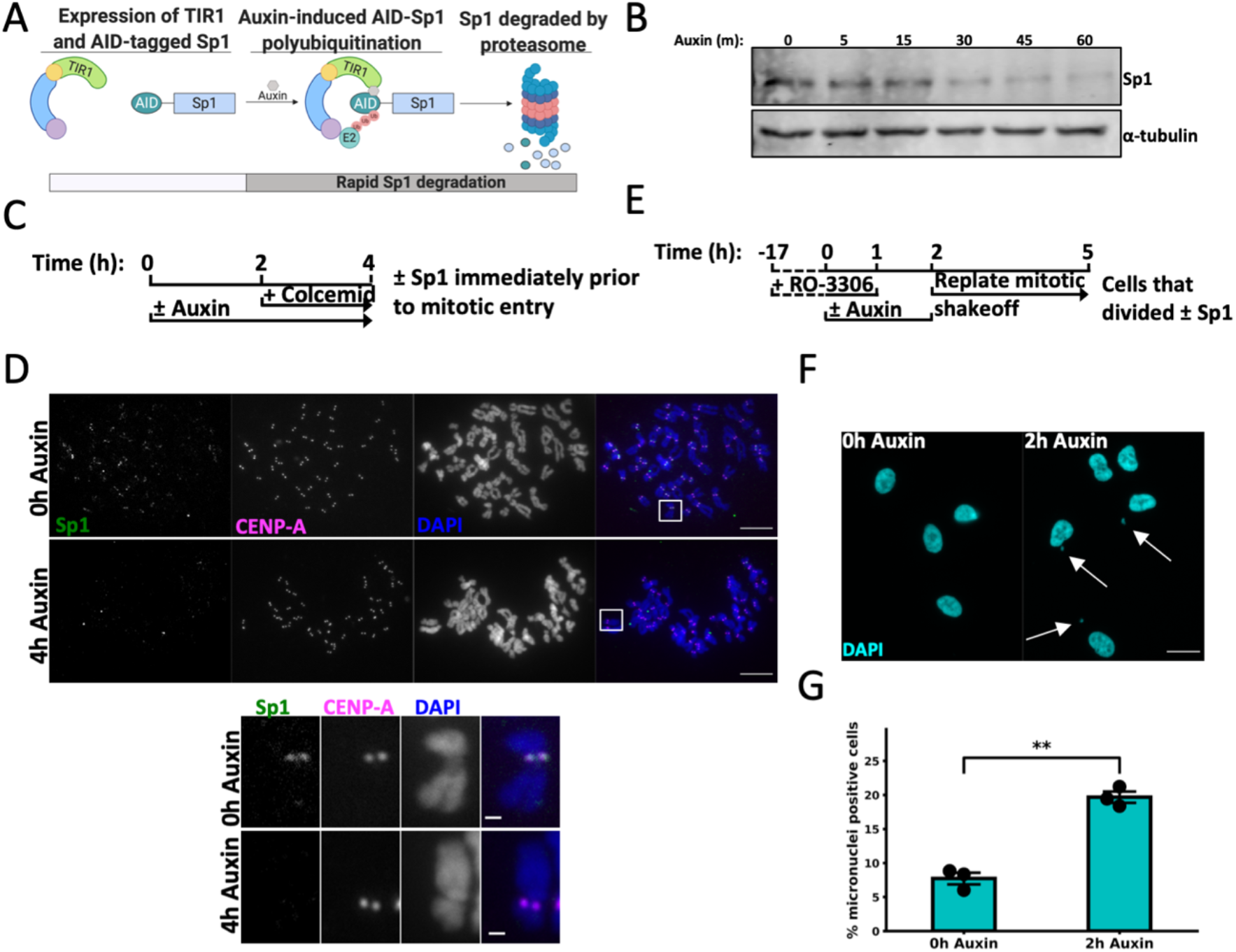
Sp1 regulates chromosome segregation during mitosis. **(A)** Schematic describing mAID-Sp1 protein degradation. Created with Biorender.com. **(B)** Immunoblot for the indicated proteins in mAID-Sp1 cells in response to 500 μM auxin. Protein lysates were collected at the indicated time points. **(C)** Schematic outlining the experimental strategy for **(D).** Upper panel: mAID-Sp1 cells were arrested in metaphase and collected following the described protocol in (Fig. 1C), spread onto a glass slide, and stained for Sp1 and CENP-A. Scale bars = 10 μm. Squares indicate the inset (Lower panel). Scale bars = 1 μm. **(E)** Schematic outlining the experimental strategy for (F). **(F)** Fluorescent detection of DAPI-stained interphase chromosomes following the indicated treatment. **(G)** The percentage of cells harboring micronuclei (white arrows). Scale bar = 10 μm. Minimum 150 cells counted per treatment. n = 3. Black circles represent the mean of each biological replicate. Error bars represent s.e.m. p = 0.00056, unpaired *t*-test‥ All images are representative.

RNAi-mediated Sp1 depletion results in chromosome-segregation errors (Astrinidis et al., 2010). We therefore hypothesized that Sp1 is functioning at mitotic centromeres to maintain mitotic fidelity. We next evaluated whether Sp1 regulates chromosome segregation during mitosis. In order to assess chromosome segregation without disrupting the mitotic spindle, we arrested mAID-Sp1 cells at G2/M by inhibiting CDK1 with 7.5 μM RO-3306 (CDK1i). We treated arrested cells with auxin for one hour then released the cells into mitosis by washing out CDK1i. One hour after release, we isolated mitotic cells by shakeoff and allowed them to enter G1. This strategy ensures that we are only evaluating cells that have undergone mitosis shortly after Sp1 was degraded (Fig. 2E). We then quantified the percentage of cells containing micronuclei, a marker of chromosome segregation errors. We determined that rapid Sp1 depletion results in a statistically significant increase in the percentage of micronuclei positive cells, indicating that Sp1 regulates chromosome segregation during mitosis (Fig. 2F, G, Supplemental Fig. 1A). Together, we show that Sp1 dynamically localizes to mitotic centromeres and regulates chromosome segregation during mitosis.

### Sp1 regulates mitotic progression

In order to gain a more comprehensive understanding of Sp1’s role during mitosis, we evaluated mitotic progression in live cells. To visualize whole chromosomes in the absence of Sp1, we depleted Sp1 in mAID-Sp1 RPE-1 cells that express H2B-mCherry (mAID-Sp1; H2B-mCherry) by treating with auxin for two hours. We then imaged these cells every three minutes for four hours (Fig. 3A, B, Supplemental Fig. 1B, and Movies 1, 2). Rapid Sp1 depletion results in several aberrant mitotic phenotypes. First, Sp1 depletion results in dramatic increases in both total mitotic duration and time from nuclear envelope breakdown (NEBD) to anaphase (Fig. 3C, D). The difference in NEBD to anaphase is similar to the difference in total mitotic duration, indicating that Sp1 is regulating mitotic progression from NEBD to anaphase. Rapid Sp1 depletion appears to result in prolonged monopolar spindle formation (Fig. 3B). Further, careful evaluation of these image sequences revealed that Sp1 fails to properly align along the metaphase plate (Fig. 3B). In order to quantify this phenotype, we arrested mAID-Sp1 cells at the G2/M checkpoint by CDKi, depleted Sp1 with auxin, and released these cells into mitosis and arrested them in metaphase by treating with 10 μm MG132 for thirty minutes after release (Fig. 3E). This strategy ensured that we evaluated the chromosome alignment specifically in metaphase cells that had progressed through mitosis without Sp1. We then quantified the percentage of these cells harboring rogue chromosomes separate from the primary cluster of chromosomes. We determined that Sp1 regulates metaphase alignment during mitosis (Fig. 3F, G).

**Fig. 3.**
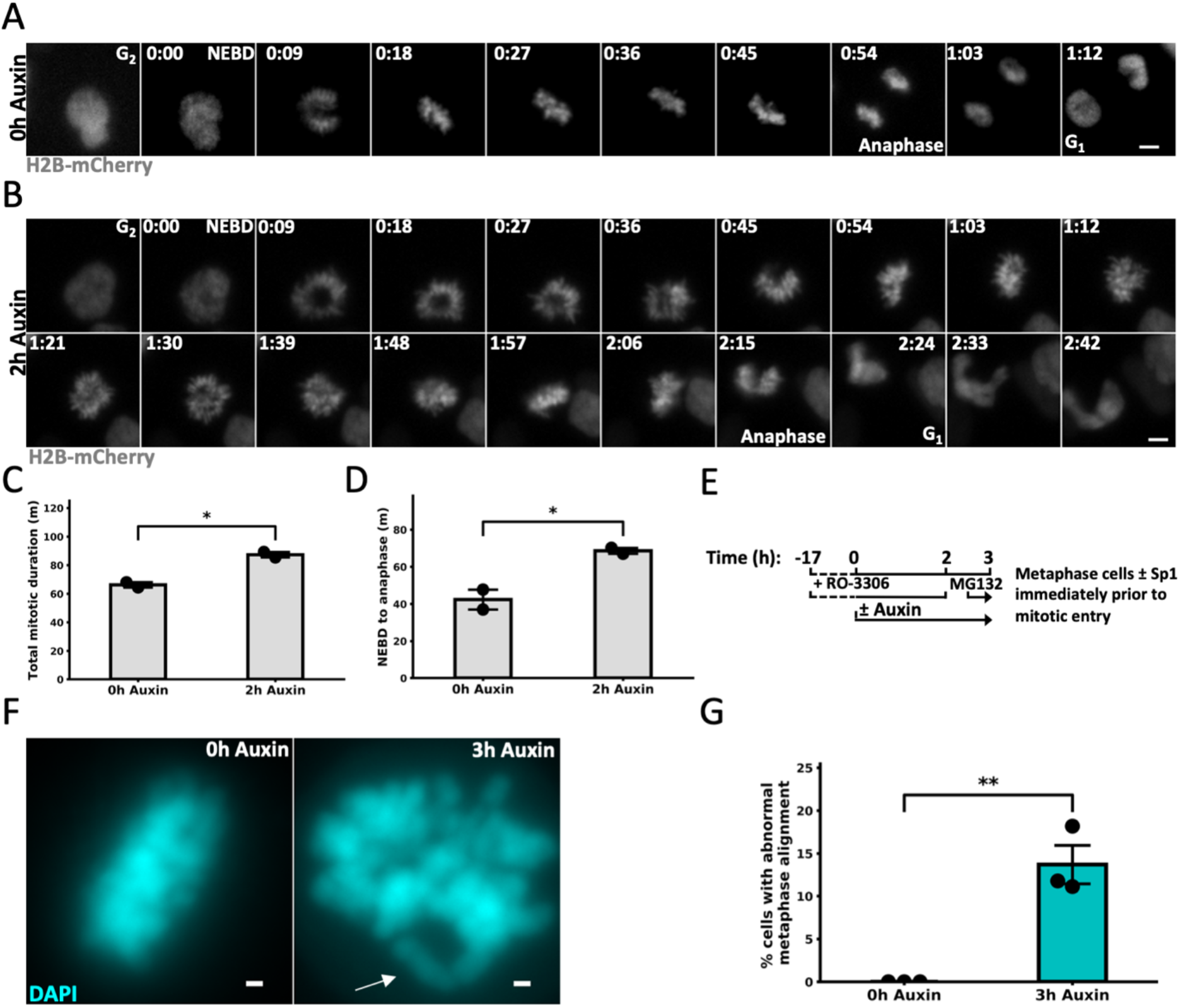
Sp1 regulates mitotic progression. **(A,B)** Live cell imaging of mAID-Sp1; H2B-mCherry cells following the indicated treatments. While images were taken every three minutes, the above image sequence represents images taken every nine minutes to best highlight the differences between the treatments. Time = h:min. Scale bar = 5 μm. **(C)** Time (m) from nuclear envelope breakdown to G_1_. 40 cells counted per treatment. n = 2. Black circles represent the mean of each biological replicate. Error bars represent s.e.m. p = 0.016, unpaired *t*-test‥ **(D)** Time (m) from nuclear envelope breakdown to anaphase. 40 cells counted per treatment. n = 2. Black circles represent the mean of each biological replicate. Error bars represent s.e.m. p = 0.042, unpaired *t*-test‥ **(E)** Schematic outlining the experimental strategy for (F). **(F)** Fluorescent detection of DAPI-stained chromosomes in mAID-Sp1 cells that were arrested in metaphase with MG132. Misaligned (white arrow) chromosomes are completely distinguishable from the metaphase plate. Scale bar = 1 μm. **(F)** Quantification of (E). Minimum 30 cells counted per treatment. n = 3. Black circles represent the mean of each biological replicate. Error bars represent s.e.m. p = 0.0037, unpaired *t*-test. All images are representative.

### Sp1 regulates mitotic chromosome assembly though condensin complex I localization

We next investigated how Sp1 regulates chromosome segregation during mitosis. Rapid Sp1 depletion results in micronuclei formation, increased mitotic duration, aberrant metaphase alignment, and monopolar spindle formation. These phenotypes are all associated with defective chromosome condensation (Hirota et al., 2004; Martin et al., 2016; Samejima et al., 2018; Samoshkin et al., 2009). We therefore hypothesized that Sp1 regulates chromosome condensation during mitosis. We first evaluated global mitotic chromosome condensation by quantifying the percentage of cells with a general condensation defect. We categorized sister chromatid pairs without clearly distinguishable p and q arms as abnormally condensed chromosomes. Following the protocol in Fig. 2C, we found that rapid depletion of Sp1 immediately prior to mitosis results in defective chromosome condensation (Fig. 4A, B, Supplemental Fig. 1C). We next considered how Sp1 is regulating chromosome condensation by evaluating the localization of condensin complexes I and II to metaphase chromosomes. We found that rapid depletion of Sp1 immediately prior to mitosis results in decreased chromosomally associated nCAP-D2, indicating a loss of condensin complex I (Fig. 4C, D). We did not observe a decrease in nCAP-H2 at mitotic chromosomes in these cells, indicating that condensin complex II localizes to mitotic cells independently of Sp1 (Fig. 4E, F). We also detected a decrease in chromosomally associated SMC4, a subunit common to both condensin complexes (Fig 4G). Importantly, Sp1 depletion does not alter condensin complex protein levels, indicating that Sp1 regulates the chromosomal localization of condensin complex I through a nontranscriptional mechanism (Fig. 4H). Together, these results suggest that Sp1 regulates condensin I-mediated mitotic chromosome condensation.

**Fig. 4.**
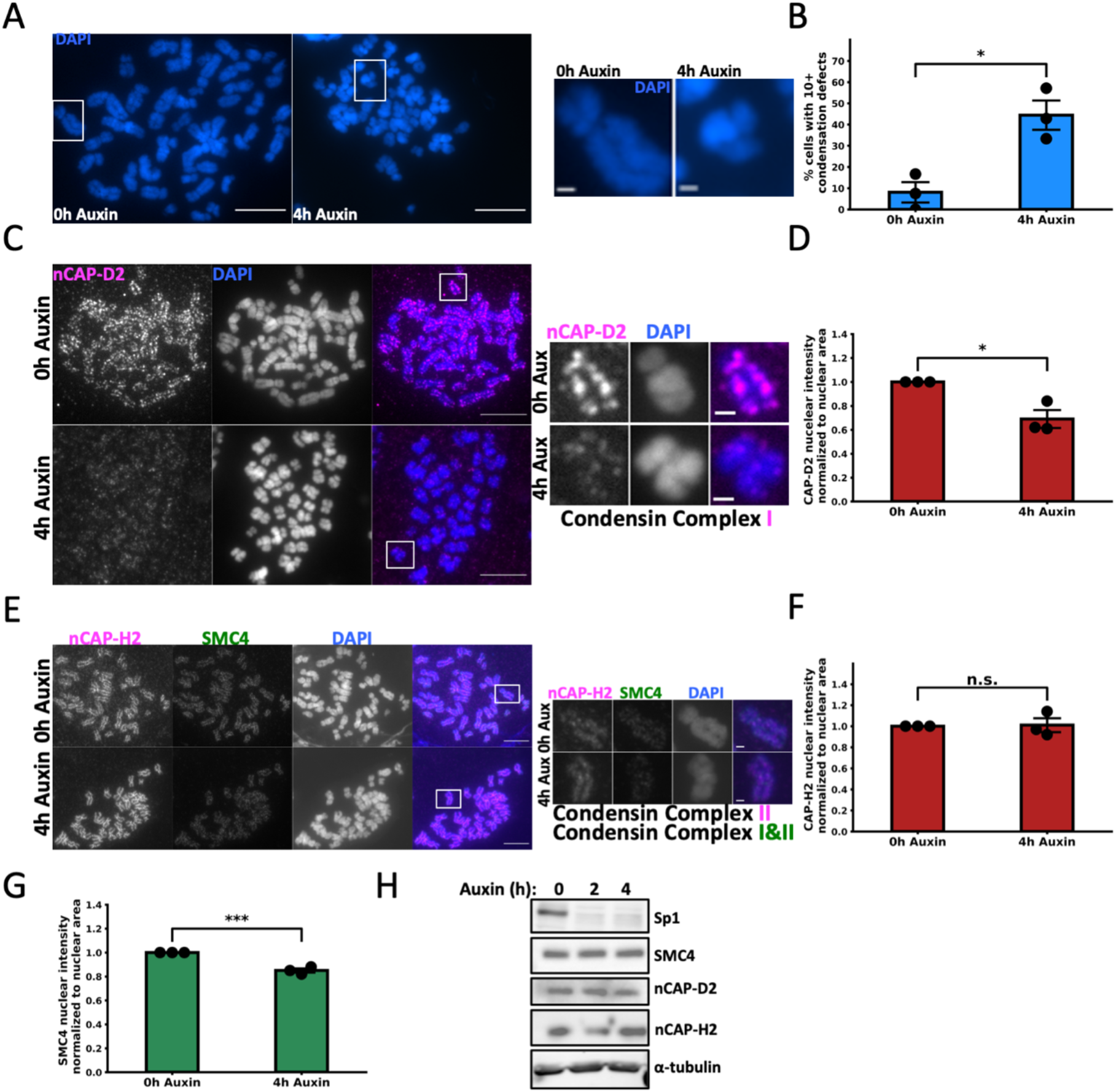
Sp1 regulates mitotic chromosome assembly though condensin complex I localization. **(A)** Left: Fluorescent detection of DAPI-stained chromosomes in mAID-Sp1 cells following the indicated treatment. mAID-Sp1 cells were arrested in metaphase, collected following the described protocol in (Fig. 2C), and spread onto a glass slide. Scale bar = 10 μm. Squares indicate the inset (Right). Scale bar = 1 μm. **(B)** Quantification of (A). Minimum 28 cells (1,288 estimated chromosomes) counted per treatment. n = 3. Black circles represent the mean of each biological replicate. Error bars represent s.e.m. p = 0.012, unpaired *t*-test‥ **(C)** Left: mAID-Sp1 cells were arrested in metaphase and collected following the described protocol in (Fig. 2C), spread onto a glass slide, and stained for CAPD-2. Scale bars = 10 μm. Squares indicate the inset (Right). Scale bar = 1 μm. **(D)** Quantification of (A). Minimum 11 cells (506 estimated chromosomes) counted per treatment. n = 3. Black circles represent the mean of each biological replicate. Error bars represent s.e.m. p = 0.015, unpaired *t*-test‥ **(E)** Left: mAID-Sp1 cells were arrested in metaphase and collected following the described protocol in (Fig. 2C), spread onto a glass slide, and stained for CAP-H2 and SMC4. Scale bars = 10 μm. Squares indicate the inset (Right). Scale bar = 1 μm. **(F)** Quantification of CAP-H2 intensity in (E). Minimum 19 cells (874 estimated chromosomes) counted per treatment. n = 3. Black circles represent the mean of each biological replicate. Error bars represent s.e.m. p = 0.89, unpaired *t*-test‥ **(H)** Quantification of SMC4 intensity in (E). Minimum 19 cells (874 estimated chromosomes) counted per treatment. n = 3. Black circles represent the mean of each biological replicate. Error bars represent s.e.m. p = 0.00098, unpaired *t*-test‥ **(H)** Representative immunoblot for the indicated proteins in mAID-Sp1 cells in response to 500 μM auxin. Protein lysates were collected at the indicated time points.

### Sp1 regulates Aurora B kinase activation early in mitosis

We next evaluated how Sp1 is regulating condensin I-mediated mitotic chromosome assembly. Aurora B kinase is required for the localization of condensin complex I but not complex II localization to mitotic chromosomes in human cells and in yeast (Lipp et al., 2007; Takemoto et al., 2007). We therefore hypothesized that Sp1 promotes Aurora B kinase activity. Aurora B kinase activity and localization varies through mitotic progression and requires an intact mitotic spindle. We controlled for this variation by first evaluating Aurora B activity in metaphase cells with an intact mitotic spindle using the protocol described in Fig. 3E. We measured Aurora B Kinase activity by immunostaining for phosphorylated INCENP (p-INCENP), a well-established marker of Aurora B activation (Salimian et al., 2011). We found that rapid Sp1 depletion prior to mitosis results in decreased p-INCENP at metaphase chromosomes (Fig. 5A, B). Aurora B activation is required for its localization to mitotic chromosomes (Salimian et al., 2011).

**Fig. 5.**
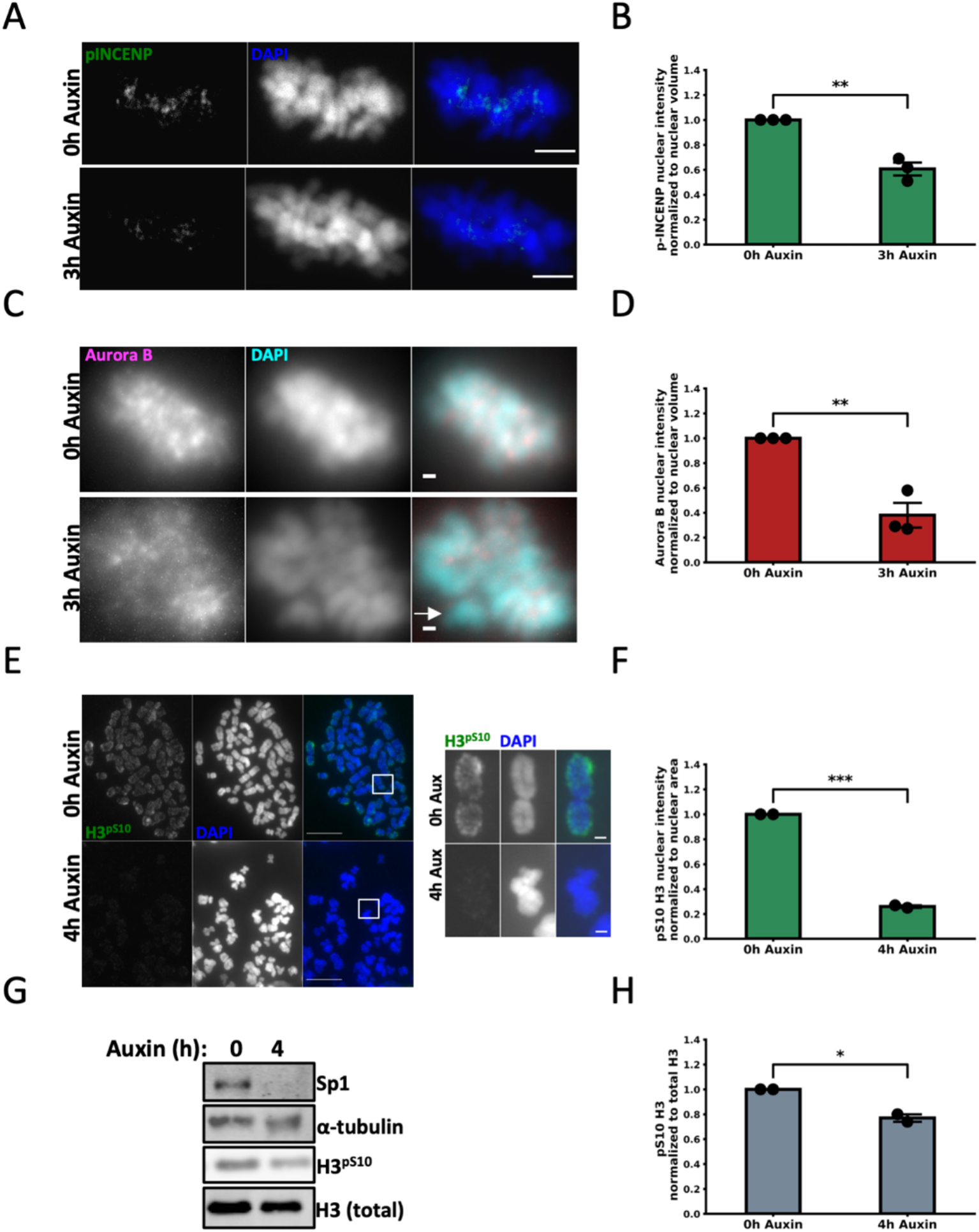
Sp1 regulates Aurora B Kinase activation early in mitosis. **(A)** mAID-Sp1 cells were arrested in metaphase following the protocol described in (Fig. 3E) and stained for p-INCENP. Scale bar = 5 μm. **(B)** Quantification of (A). Minimum 29 cells counted per treatment. n = 3. Black circles represent the mean of each biological replicate. Error bars represent s.e.m. p = 0.0017, unpaired *t*-test‥ **(C)** mAID-Sp1 cells were arrested in metaphase following the protocol described in (Fig. 3E) and stained for Aurora B kinase. Scale bar = 1 μm. White arrow indicates a misaligned chromosome. **(D)** Quantification of (C). Minimum 30 cells counted per treatment. n = 3. Black circles represent the mean of each biological replicate. Error bars represent s.e.m. p = 0.0035, unpaired *t*-test‥ **(E)** Left: mAID-Sp1 cells were arrested in metaphase and collected following the described protocol in (Fig. 2C), spread onto a glass slide, and stained for H3^pS10^. Scale bar = 10 μm. Squares indicate the inset (Right). Scale bar = 1 μm. **(F)** Quantification of (A). Minimum 19 cells (874 estimated chromosomes) counted per treatment. n = 3. Black circles represent the mean of each biological replicate. Error bars represent s.e.m. p = 0.00018, unpaired *t*-test‥ **(G)** Representative immunoblot for the indicated proteins in mAID-Sp1 cells following treatment with 500 μM auxin. Protein lysates were collected at the indicated time points. **(H)** Quantification of the densitometry normalized to H3 (total) from (G). n = 2. Black circles represent the mean of each biological replicate. Error bars represent s.e.m. p = 0.017, unpaired *t*-test.

Consistent with these observations, we next found that Sp1 regulates Aurora B kinase localization to metaphase chromosomes (Fig. 5C, D). Importantly, Aurora B is not enriched at the misaligned chromosome (Fig. 5C), indicating that Aurora B is not functioning properly in the absence of Sp1. Additionally, Aurora B is not decreased at the protein level, indicating that Sp1 regulates Aurora B localization to mitotic chromosomes through a nontranscriptional mechanism (Supplemental Fig. 1E). Aurora B kinase activity early in mitosis is responsible for condensin I localization to mitotic chromosomes. Aurora B phosphorylates histone H3 on serine 10 (H3^pS10^) during late G2/early prophase in an event linked to chromosome condensation (Crosio et al., 2002). We therefore next evaluated if Sp1 is required for this phosphorylation. Due to the stability of this mark through metaphase, we rapidly depleted Sp1 immediately prior to mitosis and immunostained metaphase cells for H3^pS10^. We found that rapid depletion of Sp1 results in a dramatic decrease in H3^pS10^, indicating that Sp1 regulates Aurora B activity during the initial phases of mitosis (Fig. 5E, F). We also observed a decrease in H3^pS10^ protein levels, corroborating our IF data (Fig. 5G, H). Together, these results indicate that Sp1 regulates Aurora B kinase activation early in mitosis.

### Sp1 expression correlates with aneuploidy in human cancers

In order to determine the clinical relevance of our findings, we examined the relationship between Sp1 and aneuploidy in human cancers by comparing Sp1 gene expression with numerical alterations to whole somatic chromosomes (Cerami et al., 2012; Gao et al., 2013; Hoadley et al., 2018; Taylor et al., 2018). Since bidirectional changes in the expression of several genes can result in chromosome segregation errors, we characterized the relationship between both decreases and increases in Sp1 expression and whole chromosome aneuploidy (WCA) (Iwanaga et al., 2007; Jeganathan, Malureanu, Baker, Abraham, & Van Deursen, 2007; Potapova & Gorbsky, 2017; Ricke, Jeganathan, & van Deursen, 2011; Sotillo et al., 2007). To control for false positives that may arise from gain or loss of the Sp1 locus on chromosome 12, we used z-scores normalized to all samples in the dataset, rather than normalized to only samples with diploid Sp1. We observed a strong negative correlation between Sp1 underexpression and whole chromosome aneuploidy (WCA) in kidney renal papillary cell carcinoma (Fig. 6A) (Schober & Schwarte, 2018). Further, we also observed a moderate negative correlation between reduced Sp1 expression and WCA in both ovarian serous cystadenocarcinoma and mesothelioma (Fig. 6B, C). These results support our findings that Sp1 is required for mitotic fidelity.

**Fig. 6.**
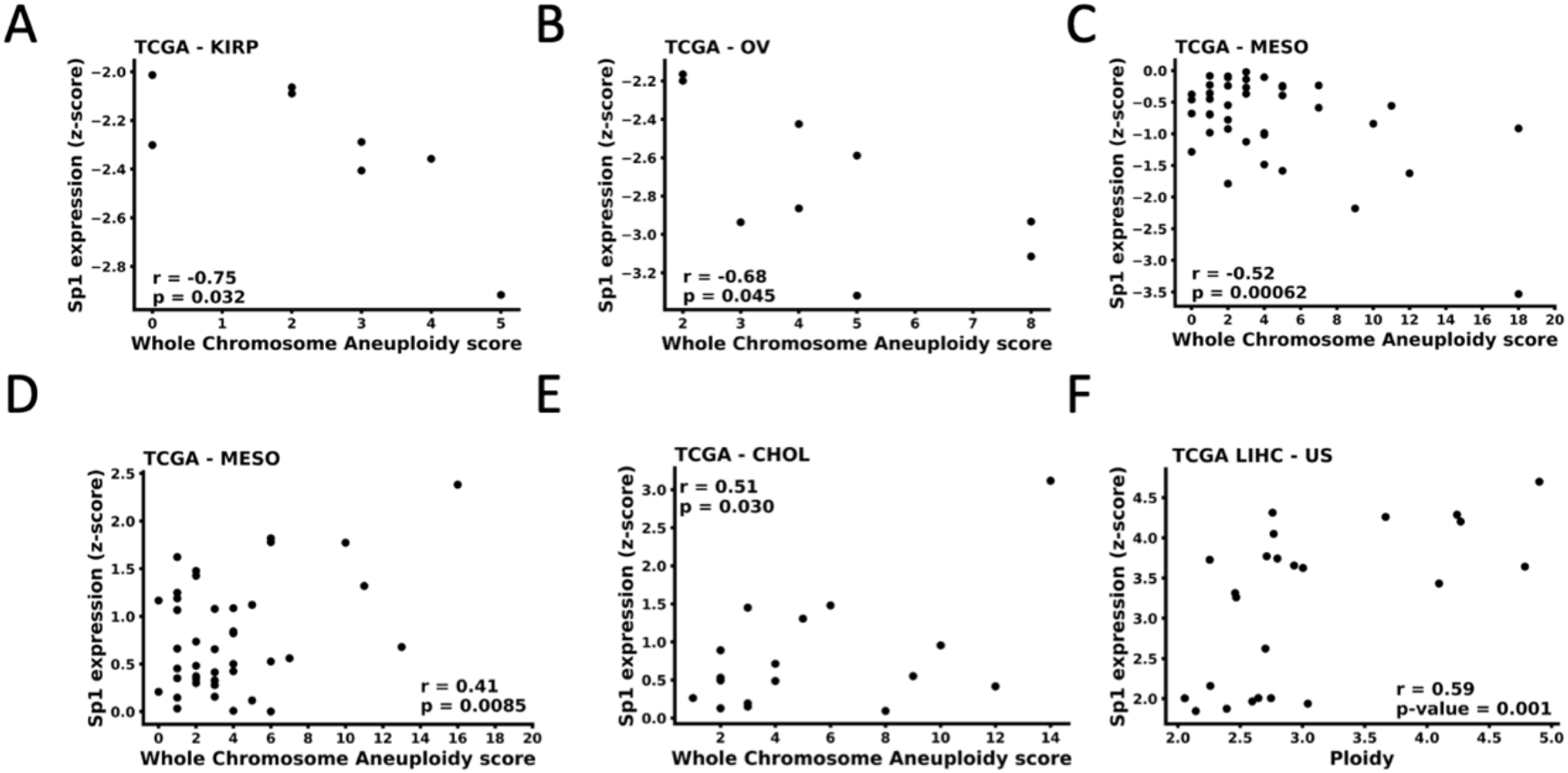
Sp1 expression correlates with aneuploidy in human cancers. Scatterplots of Sp1 expression (y-axis) and aneuploidy (x-axis) in **(A)** kidney renal papillary cell carcinoma **(B)** ovarian serous cystadenocarcinoma **(C)** and **(D)** mesothelioma **(E)** cholangiocarcinoma and **(F)** liver hepatocellular carcinoma. Correlation coefficients and p values are indicated on the graphs. For (A) and (B), Sp1 expression was not normally distributed and therefore the relationship between Sp1 expression and WCA was evaluated with Spearman correlation. All other relationships were evaluated with Pearson correlation. Negative r values indicate a negative correlation between Sp1 expression and WCA. Positive r values indicate a positive correlation between Sp1 expression and WCA.

We found that an increase in Sp1 expression also correlates with WCA in a variety of cancers. Notably, we identified a moderate positive correlation between increased Sp1 expression and WCA in mesothelioma and cholangiocarcinoma (Fig. 6D, E). We also found that Sp1 overexpression positively correlates with changes in ploidy in liver hepatocellular carcinoma (Fig 6F). These results indicate that bidirectional changes in Sp1 expression may result in a loss of mitotic fidelity. Supporting our data that Sp1 regulates chromosome segregation through aurora B signaling, bidirectional changes in Aurora B expression or activity are also associated with aneuploidy (González-Loyola et al., 2015; Kumari, Ulrich, Krause, Finkernagel, & Gaubatz, 2014; Liang et al., 2020; Ricke et al., 2011) Together, these data indicate that Sp1 has a clinically relevant relationship with aneuploidy.

## Discussion

Mitotic chromosome assembly is a complex process with broad pathophysiological implications and therefore constitutes an essential area of molecular research. The role of transcription factors in this process has been largely overlooked. The present study reveals a role for a ubiquitously expressed transcription factor in mitotic chromosome assembly and chromosome segregation. We demonstrate that Sp1 protects genomic integrity during mitosis by promoting mitotic chromosome assembly through Aurora B kinase and condensin complex I recruitment to the chromatin. This assembly is then required for proper chromosome segregation during anaphase (Fig. 7).

**Fig 7.**
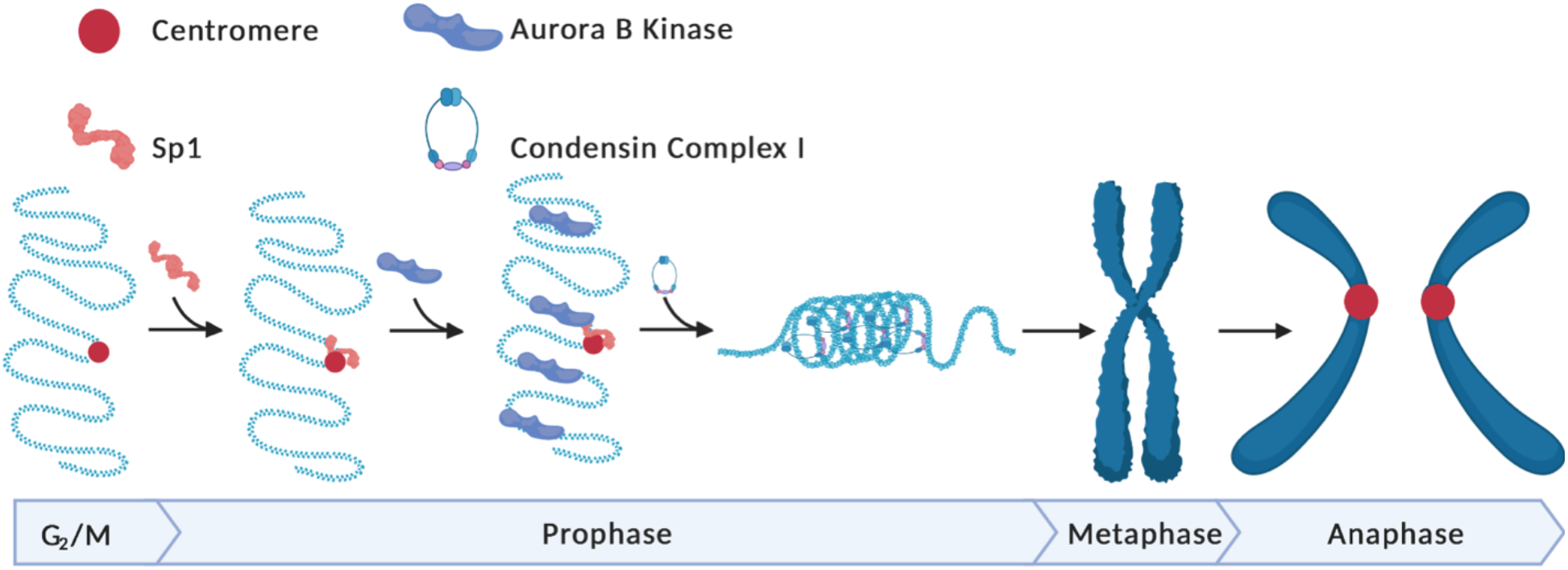
Sp1 regulates chromosome segregation during mitosis by promoting mitotic chromosome assembly Sp1 localizes to mitotic centromeres early in mitosis and regulates aurora B kinase recruitment, condensin I-mediated chromosome assembly in prophase and ultimately proper segregation of sister chromatids in anaphase. Model created with Biorender.com.

The role of transcription factors in mitotic chromosome assembly and chromosome segregation is an emerging area of research. Transcription factors have recently been shown to localize to mitotic chromatin through a highly dynamic process that may be independent of sequence-specific DNA binding (Ginno et al., 2018; Raccaud et al., 2019; Teves et al., 2016). This localization has been implicated in mitotic bookmarking, which is required to maintain transcriptional programs upon entry into G1 (Kadauke & Blobel, 2013; Liu et al., 2017; Teves et al., 2016). There is now growing support for a more direct role for transcription factors in mitosis-specific processes. In yeast, transcription factors are required for chromosome condensation and mitotic chromosome assembly (Iwasaki et al., 2015; K.-D. Kim, Tanizawa, Iwasaki, & Noma, 2016; Sutani et al., 2015). However, the role of mammalian transcription factors in these processes is unclear. Here, we implicate a mammalian transcription factor in promoting mitotic chromosome assembly.

We predict that Sp1 mediates mitotic chromosome assembly independently of its role as a transcriptional regulator. Auxin-mediated rapid Sp1 depletion results in defective chromosome assembly and segregation within approximately two hours, which is likely too transient to be a result of perturbed expression of mitotic regulators (Fig. 2 – 5). Indeed, rapid Sp1 degradation does not alter the protein levels of condensin complex factors or Aurora B kinase (Fig. 4H, Supplemental Fig. 1E). However, these data do not preclude the possibility that Sp1 regulates chromosome condensation by promoting transcription per se at regions where condensin is loaded. In yeast, condensins are loaded onto DNA that is either actively transcribed or bound by transcription factors (Iwasaki et al., 2015; K.-D. Kim et al., 2016; Sutani et al., 2015). Therefore, Sp1 may be regulating chromosome condensation by facilitating a permissive chromatin environment for condensin loading or recruiting condensin directly to Sp1 bound chromatin. However, rapid degradation of Sp1 at mitotic centromeres results in chromosome condensation defects, indicating that Sp1 is likely not functioning directly along mitotic chromosome arms to promote condensin I loading (Fig. 2D, 4). We therefore conclude that Sp1 does not promote chromosome condensation through transcriptional regulation of condensin complex protein levels or their association with mitotic chromosomes.

We next considered how Sp1 may function at mitotic centromeres to promote chromosome condensation. The catalytic subunit of the chromosome passenger complex (CPC), Aurora B kinase, is responsible for condensin complex I loading early during mitosis (Lipp et al., 2007; Takemoto et al., 2007). The CPC is recruited to the centromere in late G2 before being targeted to prophase chromosome arms to promote mitotic chromosome condensation (Abad et al., 2019; Ainsztein, Kandels-Lewis, Mackay, & Earnshaw, 1998; Seul Kim et al., 2020; Ruppert et al., 2018). Rapid Sp1 degradation results in loss of CPC activity early during mitosis (Fig. 5E-H), which accounts for the loss of mitotic chromosome assembly (Fig. 4) (Lipp et al., 2007; Takemoto et al., 2007). Consistent with our results, loss of CPC activity partially phenocopies condensin dysfunction, including chromosome segregation errors, aberrant metaphase alignment and disrupted mitotic chromosome assembly (Fig. 2F-G, Fig. 3F-E, Fig. 4) (Adams, Maiato, Earnshaw, & Carmena, 2001; Lipp et al., 2007; Martin et al., 2016; Samoshkin et al., 2009). We therefore conclude that Sp1 is regulating mitotic chromosome assembly and segregation through CPC recruitment and activation at the centromere early during mitosis (Fig. 7).

Determining how Sp1 is regulating the CPC is challenging. Although the CPC has diverse functions throughout mitotic progression, most studies focus on its role later during mitosis (Carmena et al., 2012). Therefore, little is known about how the CPC is activated and recruited early during mitosis. Emerging evidence suggests that the prophase localization of the CPC requires unique chromatin architecture at both the centromere and along chromosome arms (Abad et al., 2019; Seul Kim et al., 2020). Sp1 is known to interact with a variety of chromatin modifying factors to generate permissive chromatin environments for diverse functions including transcription and DNA repair (Beishline & Azizkhan-Clifford, 2015; Beishline, Sravanthi, & and Azizkhan-Clifford, 2013; Doetzlhofer et al., 1999; Hung, Wang, & Chang, 2006; Kadam et al., 2000). We speculate that Sp1 may be performing a similar function at mitotic centromeres by facilitating the dynamic changes in chromatin required for CPC localization. This characterization is challenging due to the paucity of information about the requirements for CPC localization early during mitosis. Another intriguing possibility is that Sp1 is regulating transcription through the centromere, which is required for Aurora B localization and function in *Xenopus* eggs (Blower, 2016; Grenfell, Heald, & Strzelecka, 2016). However, these studies have not evaluated if this transcription is occurring early or later during mitosis. In humans, centromeric transcription is initiated during late mitosis and is therefore not responsible for CPC-induced mitotic chromosome condensation during prophase (Quénet & Dalal, 2014). We therefore predict that Sp1 is not regulating CPC-dependent mitotic chromosome assembly by mediating transcription at the centromere. Overall, we implicate the transcription factor Sp1 as a key mediator for CPC localization and function early during mitosis and predict that this function is independent of Sp1’s transcriptional activity (Fig. 7).

Sp1’s mitosis-specific effects may be clinically actionable. We demonstrate here that Sp1 expression correlates with aneuploidy in human cancers (Fig. 6). Augmenting these data, Sp1 expression is dysregulated in a variety of cancers and is a poor prognostic indicator (Beishline & Azizkhan-Clifford, 2015). However, despite the pervasiveness of Sp1 dysregulation in cancer, its ubiquity and essential role in normal cells render it a poor therapeutic target. However, we may be able to indirectly target Sp1 in cancer by identifying and exploiting its synthetic lethal dependencies. Several recent studies have identified synthetic lethal interactions during mitosis in other contexts (Oser et al., 2019; van der Lelij et al., 2017). Therefore, characterizing a mitosis-specific role for Sp1 may potentiate its clinical impact. We will explore this possibility in future studies.

Defective mitotic chromosome assembly is clinically relevant. Dysregulation of condensin complex I and II are both associated with poor prognosis and are gaining attention as potential therapeutic targets (Wang, Yang, Li, & Cao, 2018). High nCAP-D2 expression is associated with increased metastasis and decreased survival in triple negative breast cancer (Y. Zhang et al., 2020). Overexpression of another condensin complex I subunit, nCAP-G, is associated with multidrug resistance in colorectal cancer and proliferation and migration in hepatocellular carcinoma (Q. Zhang, Su, Shan, Gao, & Wu, 2018). Similarly, upregulation of condensin complex I subunit nCAP-H is associated with drug resistance and decreased disease-free survival in oral squamous cell carcinoma (Shimomura, Sasahira, Nakashima, Kurihara-Shimomura, & Kirita, 2019). The aurora B kinase – condensin axis is also dysregulated in disease. In B-cell acute lymphoblastic leukemia, this axis is considered a pathogenic contributor to high-hyperdiploidy, a common and initiating oncogenic event (Molina et al., 2020). Aurora B kinase itself is a longstanding target in cancer therapy (Helfrich et al., 2016; Tang et al., 2017; Wilkinson et al., 2007). These data highlight the clinical relevance of mitotic chromosome assembly. Extensive characterization of the mechanisms governing this assembly may therefore result in a significant clinical impact in cancer diagnosis and therapy.

Here, we have demonstrated that the ubiquitously expressed transcription factor Sp1 regulates mitotic chromosome assembly. Loss of Sp1 results in loss of condensin complex I localization to mitotic chromosomes and aurora B kinase dysfunction early in mitosis. Ultimately, these defects result in aberrant mitotic progression, defective metaphase alignment, and increased chromosome segregation errors. Underscoring the clinical relevance of our work, dysregulated Sp1 expression correlates with aneuploidy in a variety of contexts. Ultimately, this work challenges the paradigm that transcription factors are mitotic spectators by implicating an understudied, yet ubiquitous, factor in a disease-relevant process.

## Materials and Methods

### Cell lines

All cells were maintained at 37°C in a humidified atmosphere with 5% CO2. hTERT RPE-1 cells (ATCC) were cultured in Dulbecco’s Modified of Eagle’s Medium/Ham’s F-12 50:50 Mix (Cellgrow) supplemented with 10% fetal bovine serum (FBS; Gemini), and 0.01 mg/ml hygromycin B (Thermo Fisher Scientific). NHDF cells (ATCC) supplemented with 10% FBS. HEK293T cells were maintained in DMEM (Cellgrow) supplemented with 10% heat-inactivated FBS and Pen-Strep. HEK293-GPG were cultured in DMEM (Cellgrow) supplemented with 10% heat-inactivated FBS, Pen-Strep, 1 μg/mL tetracycline, 2 μg/mL puromycin, and 0.3 mg/mL G418. mAID-Sp1 cells were derived by first transducing RPE-1 cells with lentivirus containing sgSp1 and with lentivirus containing mAID-Sp1 downstream of the Sp1 promoter. These cells were then colony selected and screened for mAID-Sp1 expression. mAID-Sp1 expressing cells were then transduced with retrovirus containing osTIR1, challenged with Blasticidin, colony selected, and then screened for rapid depletion of Sp1 in response to auxin treatment. mAID-Sp1; H2B-mCherry cells were derived by transducing mAID-Sp1 cells with lentivirus containing mCherry-H2B.

### Drug Treatments

Cells were treated with auxin (Abcam ab146403, dissolved in ddH2O), RO-3306 (Selleck 7747, dissolved in DMSO), MG132 (Sigma 10012628, dissolved in DMSO, Colcemid (Sigma, dissolved in DMEM), and Biotin (Sigma 29129, dissolved in DMEM). All negative controls were treated with the equivalent volume of solvent. Note: in ure 4A we treated with MG132 immediately following RO-3306 washout. Because MG132 blocks the auxin-inducible proteasomal degradation of Sp1, degradation of Sp1 was not as effective (evident in Supplemental Fig. 1D). We therefore waited 30 minutes after RO-3306 washout for subsequent experiments (Fig. 3E and 5C). This waiting period appeared to facilitate Sp1 depletion (Supplemental Fig. 1E).

### Plasmids

Sp1 sgRNA constructs were made using lentiCRISPR v2,s a gift from Feng Zhang (Addgene 49535) (Samejima et al., 2018). Plasmid was cut using Bsmb1 and ligated to oligomers with the following sequence for Sp1: Forward (5’ CAC CGC ATG GAT GAA ATG ACA GCT G 3’) and Reverse (5’ AAA CCA GCT GTC ATT TCA TCC ATG C 3’) and non-targeting: Forward(5’ CAC CGG AGC CCG ACT AAA GAG GCC G 3’) and Reverse (5’ AAA CCG GCC TCT TTA GTC GGG CTC C 3’). Flag-tag in lentiCRISPr v2 with sgRNA constructs was deleted by excising Flag-Cas9 using restriction enzymes Age1 and BamH1. mAID-Sp1 was constructed by cloning the miniAID protein sequence upstream of sgSp1-resistant Sp1 protein sequence into the pLZS-Sp1 vector using Gibson assembly (Sanjana, Shalem, & Zhang, 2014). miniAid was cloned from pcDNA5/FRT EGFP-miniAID, a gift from Andrew Holland (Addgene plasmid # 101714). pLZS was generated by performing Gibson assembly to replace the CMV promoter with the endogenous Sp1 core promoter (−1612 to +1) in the pLENTI CMV GFP Zeo vector, a gift from Eric Campeau and Paul Kaufman (Addgene plasmid #17449) (Gibson et al., 2009) . pCDNA5/FRT EGFP-miniAID was a gift from Andrew Holland (Addgene plasmid # 101714). pBabe Blast osTIR1-9myc was a gift from Andrew Holland (Addgene plasmid #80073). pLENTI H2B-mCherry was generated by replacing GFP in the pLENTI CMV GFP NEO vector with H2B-mCherry using Gibson assembly. H2B-mCherry was a gift from Robert Benezra (Addgene plasmid #20972) and pLENTI CMV GFP NEO was a gift from Eric Campeau and Paul Kaufman (Addgene plasmid #17447) (Campeau et al., 2009) pBabe Blast osTIR1-9myc was a gift from Andrew Holland (Addgene plasmid #80073). All primer sequences are available upon request.

### Indirect Immunofluorescence

Cells were arrested in metaphase with 100 ng/mL colcemid (Sigma) for two hours and collected by mitotic shake off. Pelleted cells were then resuspended in hypotonic solution (10 mM Hepes pH 7.3; 2% FBS; 30 mM Glycerol; 1.0 mM CaCl2; 0.8 mM MgCl2) and incubated at 4°C for 15 minutes. Swollen cells were then spun onto a glass microscope slide using a Shandon Cytospin 3 (2000 RPM for 20 minutes). Cells were then fixed in methanol (−20°C for 30 minutes) and acetone (−20°C for 30 seconds) and allowed to dry at room temperature. Cells then were stored at −20°C indefinitely. Alternatively, cells were seeded onto coverslips and grown to ~80% confluence. Cells were then washed with PBS, fixed in 4% formaldehyde for 10 minutes, washed with PBS, permeabilized in PBS + 0.5% Triton X-100, washed in PBST (PBS + 0.1% Tween-20), blocked overnight in PBST + 3% BSA. Both coverslip-bound cells and metaphase spreads were then incubated (37°C for 30 minutes) with the following antibodies: Sp1 (1:200, Santa Cruz sc14027), CENP-A (1:200, Abcam ab13939), nCAP-D2 (1:50, Santa Cruz sc-398850), nCAP-H2 (1:50, Santa Cruz, sc-393333), SMC4 (1:1000, Novus NBP1-86635), p-INCENP (1:100, kind gift from Dr. Ben Black (University of Pennsylvania)), Aurora B Kinase (1:1000, BD 611082) and Histone H3^pS10^ (1:100, Cell Signaling #9701s). Cells were then washed with PBST and incubated (37°C for 30 minutes) with the following secondary antibodies: α-Rabbit IgG (H+L) Alexa Fluor® 488 conjugate (A21206, 1:1000), α-Mouse IgG (H+L) Alexa Fluor® 594 conjugate (A21203, 1:1000). DNA was stained (37°C for 5 minutes) with 1μg/ml DAPI (Sigma) in PBS. Cells were then washed with PBST, PBS, and H2O before being mounted in VectaMount AQ (Vector Laboratories, Inc) and stored at 4°C indefinitely. Images for Fig. 1D, 2D, 2F were obtained with the Olympus AX-70 compound microscope and iVision Scientific Image Process software by BioVision Technologies. Images for Fig. 5A were obtained with the Olympus FV3000 confocal microscope. All other images were obtained with the Evos FL compound microscope. All paired images were acquired in parallel using the same microscope settings for channels that are compared in the analysis (e.g. nCAP-D2 in 0h auxin and 4h auxin cells).

### Image Analysis

All images were processed and analyzed in parallel using Image J. Positional intensities (Fig. 1C) were obtained by drawing a line through each centromere pair, straightening the image, and deriving the positional intensities of CENP-A and Sp1 using the plot profile function in Image J. These intensities were background corrected by subtracting the first measured intensity outside of the centromere pair from all reported values. Metaphase spread images were analyzed with the following workflow: define nuclear area by converting thresholded DAPI staining to a mask; quantify that area by adding the total number of pixels in the mask; quantify chromosome-associated antibody of interest (AoI) intensity by multiplying the AoI by the DAPI mask; normalize by dividing chromosome-associated AoI intensity by total nuclear area. Aurora B Kinase images were analyzed with the following workflow: define nuclear area by converting thresholded DAPI staining to a mask; quantify that area by adding the total number of pixels in the mask; quantify nuclear Aurora B Kinase intensity by multiplying Aurora B Kinase intensity by the DAPI mask; obtain the volume of these measurements by adding together all nuclear Aurora B Kinase intensities in each z-stack slice. Due to high background, the p-INCENP images were analyzed with the following workflow: define nuclear area by converting thresholded DAPI staining to a mask; quantify that area by adding the total number of pixels in the mask.; quantify nuclear p-INCENP by summing the product of p-INCENP intensity and the DAPI mask. Determine non-nuclear p-INCENP intensity (NNI) by subtracting nuclear p-INCENP intensity from total p-INCENP intensity. Determine background intensity/pixel by dividing total NNI by total number of NNI pixels. Determine nuclear background intensity by multiplying background intensity/pixel by nuclear area. Background correct the nuclear p-INCENP by subtracting nuclear background intensity from nuclear p-INCENP intensity. Obtain the volume of these measurements by adding the background corrected nuclear p-INCENP intensity for each slice in the z-stack. Note that rapid SMC2 depletion does not change the total nuclear volume, indicating that normalizing p-INCENP signal to nuclear volume would not be affected by chromosome condensation defects (Samejima et al., 2018).

### Live Cell Imaging

mAID-Sp1; H2B-mCherry cells were grown in 8 well Falcon Chambered Cull Culture Slides (Corning 354108) for at least 24 hours. Two hours prior to imaging, cells were treated with 500 μm auxin. Images were acquired at 20x magnification every three minutes using the Evos FL auto microscope with the on-stage incubator maintaining 37°C).

### Immunoblots

Cells were lysed in 2x SDS sample buffer (12.5 mM Tris pH 6.8; 20% glycerol; 4% SDS) and boiled. Protein concentration was determined using the Pierce BCA Protein Assay Kit (ThermoFisher 23225). Samples were then supplemented with 5% β-mercaptoethanol, boiled and vortexed. Either 8 or 10 μg of sample were used for subsequent analysis. Proteins were resolved by SDS-PAGE, transferred to polyvinylidene difluoride (PVDF) membrane; the following antibodies were diluted in TBST + 5% BSA: Sp1 Ab581 (1:1000), α-tubulin (1:1000, Cell Signaling #2244), nCAP-D2 (1:1000, Santa Cruz sc-398850), nCAP-H2 (1:1000, Santa Cruz, sc-393333), SMC4 (1:1000, Novus NBP1-86635), Aurora B Kinase (1:1000, BD 611082), Histone H3^pS10^ (1:1000, Cell Signaling #9701s), Histone H3 (1:1000, Cell Signaling #3638). Primary antibodies were then recognized by the appropriate secondary antibodies: IRDye 680RD α-Rabbit (1:10,000, Li-COR), IRDye 800CW α-Mouse (1:5000, Li-COR), HRP α-Rabbit (1:2000, Jackson ImmunoResearch Laboratories 711-036-152) and HRP α-Mouse (1:2000, Jackson ImmunoResearch Laboratories 715-056-150). nCAP-D2 and nCAP-H2 were detected with the appropriate HRP-conjugated secondary antibodies; all other primary antibodies were detected with the appropriate IR-conjugated secondary antibodies. IR-conjugated secondary antibodies were detected by the Odyssey imaging system (Li-COR) and HRP-conjugated secondary antibodies were detected by the GeneSys G:Box F3 gel imaging system (Syngene).

### TCGA Datamining

Sp1 mRNA expression data was generated by the PanCancer Atlas and downloaded from the cBioPortal (www.cBioPortal.com) (Cerami et al., 2012; Gao et al., 2013; Hoadley et al., 2018). Whole chromosome aneuploidy was calculated by adding the absolute value of the aneuploidy score (AS) for chromosomes with p and q arms that had the same AS (Taylor et al., 2018). For acrocentric chromosomes, the AS was considered whole chromosome aneuploidy. Sp1 expression data and ploidy scores in LIHC were obtained using the catalog of somatic mutations in cancer (COSMIC) (cancer.sange.ac.uk) (Tate et al., 2019).

### Statistical Analysis

All statistical analysis, data handling, and data visualization was performed with Python 3.7.7 (Python Software Foundation) using the following packages: SciPy (1.4.1), pandas (1.0.3), NumPy (1.18.1), Matplotlib (3.1.3), and iPython (7.13.0) (Hunter, 2007; McKinney, 2010; Oliphant, 2006; Perez & Granger, 2007; van der Walt, Colbert, & Varoquaux, 2011; Virtanen et al., 2020). TCGA data distribution was assessed with the Shapiro-Wilk test. Because Sp1 z-scores were not normally distributed in the KIRP and OV datasets, the relationship between Sp1 expression and WCA was evaluated with Spearman’s correlation. Alternatively, since both Sp1 z-scores and ploidy were normally distributed in all other datasets, their relationship was evaluated using Pearson’s correlation. All correlation coefficients were interpreted as described (Schober & Schwarte, 2018). All other p-values were calculated with the unpaired *t*-test.

## Supporting information

Supplemental Table - list of primer sequences

## Acknowledgements

We would like to thank Dr. Shae Padrick for help with image analysis and Kelly Donovan for project advice and help with cloning. We would also like to thank all other members of the Clifford Lab for intellectual input and support.

---

**Fig. S1.**
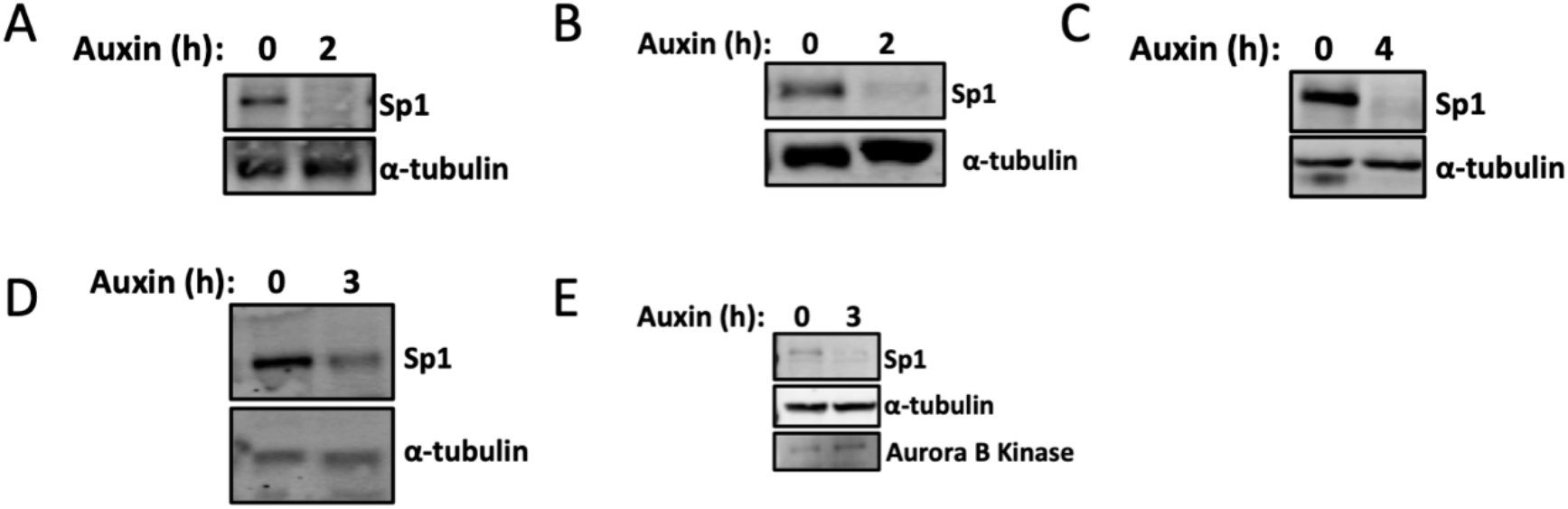
Western blots related to fig. 2-5. Representative immunoblots from **(A)** Fig. 2 D, E **(B)** Fig. 3 A-D **(C)** Fig. 4 A-G, **(D)** Fig. 5 A, B and **(E)** Fig. 3 E, F and Fig. 5 C, D.

**Video 1 and 2.** Sp1 regulates mitotic progression.

Live cell imaging of mAID-Sp1; H2B-mCherry cells following treatment with ddH2O (Video 1) or 500 μM Auxin (Video 2). Images were taken every 3 minutes. Time = h:min. Each video starts nine minutes prior to entry into mitosis. Related to Fig. 3.

## References

Abad, M. A., Ruppert, J. G., Buzuk, L., Wear, M., Zou, J., Webb, K. M., … Jeyaprakash, A. A. (2019). Borealin–nucleosome interaction secures chromosome association of the chromosomal passenger complex. Journal of Cell Biology, 218(12), 3912–3925. https://doi.org/10.1083/jcb.201905040

Adams, R. R., Maiato, H., Earnshaw, W. C., & Carmena, M. (2001). Essential roles of Drosophila inner centromere protein (INCENP) and aurora B in histone H3 phosphorylation, metaphase chromosome alignment, kinetochore disjunction, and chromosome segregation. The Journal of Cell Biology, 153(4), 865–880. https://doi.org/10.1083/jcb.153.4.865

Ainsztein, A. M., Kandels-Lewis, S. E., Mackay, A. M., & Earnshaw, W. C. (1998). INCENP Centromere and Spindle Targeting: Identification of Essential Conserved Motifs and Involvement of Heterochromatin Protein HP1. Journal of Cell Biology, 143(7), 1763–1774. https://doi.org/10.1083/jcb.143.7.1763

Astrinidis, A., Kim, J., Kelly, C. M., Olofsson, B. A., Torabi, B., Sorokina, E. M., & Azizkhan-Clifford, J. (2010). The transcription factor SP1 regulates centriole function and chromosomal stability through a functional interaction with the mammalian target of rapamycin/raptor complex. Genes Chromosomes and Cancer, 49(3), 282–297. https://doi.org/10.1002/gcc.20739

Bakhoum, S. F., & Landau, D. A. (2017). Chromosomal instability as a driver of tumor heterogeneity and evolution. Cold Spring Harbor Perspectives in Medicine, 7(6). https://doi.org/10.1101/cshperspect.a029611

Beishline, K., & Azizkhan-Clifford, J. (2015). Sp1 and the ‘hallmarks of cancer”.’ FEBS Journal, 282(2), 224–258. https://doi.org/10.1111/febs.13148

Beishline, K., Kelly, C. M., Olofsson, B. A., Koduri, S., Emrich, J., Greenberg, R. A., & Azizkhan-Clifford, J. (2012). Sp1 facilitates DNA double-strand break repair through a nontranscriptional mechanism. Molecular and Cellular Biology, 32(18), 3790–3799. https://doi.org/10.1128/MCB.00049-12

Beishline, K., Sravanthi, K., & and Azizkhan-Clifford, J. (2013). Transcription Factor Sp1 Promotes Chromatin Remodeling at DNA Double-Strand Breaks. FASEB Journal, 969–7.

Blower, M. D. (2016). Centromeric Transcription Regulates Aurora-B Localization and Activation. Cell Reports (Vol. 15). https://doi.org/10.1016/j.celrep.2016.04.054

Booth, D. G., Beckett, A. J., Molina, O., Samejima, I., Masumoto, H., Kouprina, N., … Earnshaw, W. C. (2016). 3D-CLEM Reveals that a Major Portion of Mitotic Chromosomes Is Not Chromatin. Molecular Cell, 64(4), 790–802. https://doi.org/10.1016/J.MOLCEL.2016.10.009

Campeau, E., Ruhl, V. E., Rodier, F., Smith, C. L., Rahmberg, B. L., Fuss, J. O., … Kaufman, P. D. (2009). A Versatile Viral System for Expression and Depletion of Proteins in Mammalian Cells. PLoS ONE, 4(8), e6529. https://doi.org/10.1371/journal.pone.0006529

Caravaca, J. M., Donahue, G., Becker, J. S., He, X., Vinson, C., & Zaret, K. S. (2013). Bookmarking by specific and nonspecific binding of FoxA1 pioneer factor to mitotic chromosomes. Genes & Development, 27(3), 251–260. https://doi.org/10.1101/gad.206458.112

Carmena, M., Wheelock, M., Funabiki, H., & Earnshaw, W. C. (2012). The chromosomal passenger complex (CPC): from easy rider to the godfather of mitosis. Nature Reviews Molecular Cell Biology, 13(12), 789–803. https://doi.org/10.1038/nrm3474

Cerami, E., Gao, J., Dogrusoz, U., Gross, B. E., Sumer, S. O., Aksoy, B. A., … Schultz, N. (2012). The cBio Cancer Genomics Portal: An Open Platform for Exploring Multidimensional Cancer Genomics Data: Figure 1. Cancer Discovery, 2(5), 401–404. https://doi.org/10.1158/2159-8290.CD-12-0095

Clift, D., Mcewan, W. A., Labzin, L. I., Konieczny, V., Mogessie, B., James, L. C., … Schuh, M. (2017). A Method for the Acute and Rapid Degradation of Endogenous Proteins Resource A Method for the Acute and Rapid Degradation of Endogenous Proteins. Cell, 172, 1–15. https://doi.org/10.1016/j.cell.2017.10.033

Crosio, C., Maria Fimia, G., Loury, R., Kimura, M., Okano, Y., Zhou, H., … Sassone-Corsi, P. (2002). Mitotic Phosphorylation of Histone H3: Spatio-Temporal Regulation by Mammalian Aurora Kinases. MOLECULAR AND CELLULAR BIOLOGY, 22(3), 874–885. https://doi.org/10.1128/MCB.22.3.874-885.2002

Doetzlhofer, A., Rotheneder, H., Lagger, G., Koranda, M., Kurtev, V., Brosch, G., … Seiser, C. (1999). Histone deacetylase 1 can repress transcription by binding to Sp1. Molecular and Cellular Biology, 19(8), 5504–5511. https://doi.org/10.1128/mcb.19.8.5504

Gao, J., Aksoy, B. A., Dogrusoz, U., Dresdner, G., Gross, B., Sumer, S. O., … Schultz, N. (2013). Integrative analysis of complex cancer genomics and clinical profiles using the cBioPortal. Science Signaling, 6(269), pl1. https://doi.org/10.1126/scisignal.2004088

Gerlich, D., Hirota, T., Koch, B., Peters, J.-M., & Ellenberg, J. (2006). Condensin I Stabilizes Chromosomes Mechanically through a Dynamic Interaction in Live Cells. Current Biology, 16(4), 333–344. https://doi.org/10.1016/J.CUB.2005.12.040

Gibson, D. G., Young, L., Chuang, R.-Y., Venter, J. C., Hutchison, C. A., & Smith, H. O. (2009). Enzymatic assembly of DNA molecules up to several hundred kilobases. Nature Methods, 6(5), 343–345. https://doi.org/10.1038/nmeth.1318

Ginno, P. A., Burger, L., Seebacher, J., Iesmantavicius, V., & Schübeler, D. (2018). Cell cycle-resolved chromatin proteomics reveals the extent of mitotic preservation of the genomic regulatory landscape. Nature Communications, 9(1), 4048. https://doi.org/10.1038/s41467-018-06007-5

González-Loyola, A., Fernández-Miranda, G., Trakala, M., Partida, D., Samejima, K., Ogawa, H., … Malumbres, M. (2015). Aurora B Overexpression Causes Aneuploidy and p21 Cip1 Repression during Tumor Development. https://doi.org/10.1128/MCB.01286-14

Grenfell, A. W., Heald, R., & Strzelecka, M. (2016). Mitotic noncoding RNA processing promotes kinetochore and spindle assembly in Xenopus. Journal of Cell Biology, 214(2), 133–141. https://doi.org/10.1083/jcb.201604029

Helfrich, B. A., Kim, J., Gao, D., Chan, D. C., Zhang, Z., Tan, A.-C., & Bunn, P. A. (2016). Barasertib (AZD1152), a Small Molecule Aurora B Inhibitor, Inhibits the Growth of SCLC Cell Lines In Vitro and In Vivo. Molecular Cancer Therapeutics, 15(10), 2314–2322. https://doi.org/10.1158/1535-7163.MCT-16-0298

Hirano, T. (2012). Condensins: Universal organizers of chromosomes with diverse functions. Genes and Development. https://doi.org/10.1101/gad.194746.112

Hirano, T. (2016). Condensin-Based Chromosome Organization from Bacteria to Vertebrates. Cell, 164(5), 847–857. https://doi.org/10.1016/j.cell.2016.01.033

Hirota, T., Gerlich, D., Koch, B., Ellenberg, J., & Peters, J.-M. (2004). Distinct functions of condensin I and II in mitotic chromosome assembly. Journal of Cell Science, 117, 6435–6445. https://doi.org/10.1242/jcs.01604

Hoadley, K. A., Yau, C., Hinoue, T., Wolf, D. M., Lazar, A. J., Drill, E., … Laird, P. W. (2018). Cell-of-Origin Patterns Dominate the Molecular Classification of 10,000 Tumors from 33 Types of Cancer. Cell, 173(2), 291–304.e6. https://doi.org/10.1016/j.cell.2018.03.022

Holland, A. J., Fachinetti, D., Han, J. S., & Cleveland, D. W. (2012). Inducible, reversible system for the rapid and complete degradation of proteins in mammalian cells. Proceedings of the National Academy of Sciences, 109(49), E3350–E3357. https://doi.org/10.1073/pnas.1216880109

Hung, J.-J., Wang, Y.-T., & Chang, W.-C. (2006). Sp1 deacetylation induced by phorbol ester recruits p300 to activate 12(S)-lipoxygenase gene transcription. Molecular and Cellular Biology, 26(5), 1770–1785. https://doi.org/10.1128/MCB.26.5.1770-1785.2006

Hunter, J. D. (2007). Matplotlib: A 2D Graphics Environment. Computing in Science & Engineering, 9(3), 90–95. https://doi.org/10.1109/MCSE.2007.55

Ishak, C. A., Coschi, C. H., Roes, M. V., & Dick, F. A. (2017). Cell Cycle Disruption of CDK-resistant chromatin association by pRB causes DNA damage, mitotic errors, and reduces Condensin II recruitment. Cell Cycle, 16(15), 1430–1439. https://doi.org/10.1080/15384101.2017.1338984

Iwanaga, Y., Chi, Y. H., Miyazato, A., Sheleg, S., Haller, K., Peloponese, J. M., … Jeang, K. T. (2007). Heterozygous deletion of mitotic arrest-deficient protein 1 (MAD1) increases the incidence of tumors in mice. Cancer Research, 67(1), 160–166. https://doi.org/10.1158/0008-5472.CAN-06-3326

Iwasaki, O., Tanizawa, H., Kim, K.-D., Yokoyama, Y., Corcoran, C. J., Tanaka, A., … Noma, K. (2015). Interaction between TBP and Condensin Drives the Organization and Faithful Segregation of Mitotic Chromosomes. Molecular Cell, 59(5), 755–767. https://doi.org/10.1016/J.MOLCEL.2015.07.007

Jeganathan, K., Malureanu, L., Baker, D. J., Abraham, S. C., & Van Deursen, J. M. (2007). Bub1 mediates cell death in response to chromosome missegregation and acts to suppress spontaneous tumorigenesis. Journal of Cell Biology, 179(2), 255–267. https://doi.org/10.1083/jcb.200706015

Kadam, S., McAlpine, G. S., Phelan, M. L., Kingston, R. E., Jones, K. A., & Emerson, B. M. (2000). Functional selectivity of recombinant mammalian SWI/SNF subunits. Genes & Development, 14(19), 2441–2451. https://doi.org/10.1101/gad.828000

Kadauke, S., & Blobel, G. A. (2013). Mitotic bookmarking by transcription factors. Epigenetics & Chromatin, 6(1), 6. https://doi.org/10.1186/1756-8935-6-6

Kadauke, S., Udugama, M. I., Pawlicki, J. M., Achtman, J. C., Jain, D. P., Cheng, Y., … Blobel, G. A. (2012). Tissue-specific mitotic bookmarking by hematopoietic transcription factor GATA1. Cell, 150(4), 725–737. https://doi.org/10.1016/j.cell.2012.06.038

Kim, K.-D., Tanizawa, H., Iwasaki, O., & Noma, K. (2016). Transcription factors mediate condensin recruitment and global chromosomal organization in fission yeast. Nature Genetics, 48(10), 1242–1252. https://doi.org/10.1038/ng.3647

Kim, Seul, Kim, N. H., Park, J. E., Hwang, J. W., Myung, N., Hwang, K.-T., … Kim, Y. K. (2020). PRMT6-mediated H3R2me2a guides Aurora B to chromosome arms for proper chromosome segregation. Nature Communications, 11(1), 612. https://doi.org/10.1038/s41467-020-14511-w

Kim, Seungsoo, & Shendure, J. (2019). Mechanisms of Interplay between Transcription Factors and the 3D Genome. Molecular Cell, 76(2), 306–319. https://doi.org/10.1016/J.MOLCEL.2019.08.010

Kumari, G., Ulrich, T., Krause, M., Finkernagel, F., & Gaubatz, S. (2014). Induction of p21CIP1 Protein and Cell Cycle Arrest after Inhibition of Aurora B Kinase Is Attributed to Aneuploidy and Reactive Oxygen Species. Journal of Biological Chemistry, 289(23), 16072–16084. https://doi.org/10.1074/JBC.M114.555060

Lee, A. J. X., Endesfelder, D., Rowan, A. J., Walther, A., Birkbak, N. J., Futreal, P. A., … Swanton, C. (2011). Chromosomal instability confers intrinsic multidrug resistance. Cancer Research, 71(5), 1858–1870. https://doi.org/10.1158/0008-5472.CAN-10-3604

Lee, H.-S., Lin, Z., Chae, S., Yoo, Y.-S., Kim, B.-G., Lee, Y., … Cho, H. (2018). The chromatin remodeler RSF1 controls centromeric histone modifications to coordinate chromosome segregation. Nature Communications, 9(1), 3848. https://doi.org/10.1038/s41467-018-06377-w

Liang, C., Zhang, Z., Chen, Q., Yan, H., Zhang, M., Zhou, L., … Wang, F. (2020). Centromere-localized Aurora B kinase is required for the fidelity of chromosome segregation. The Journal of Cell Biology, 219(2). https://doi.org/10.1083/jcb.201907092

Lipp, J. J., Hirota, T., Poser, I., & Peters, J. M. (2007). Aurora B controls the association of condensin I but not condensin II with mitotic chromosomes. Journal of Cell Science, 120(7), 1245–1255. https://doi.org/10.1242/jcs.03425

Liu, Y., Pelham-Webb, B., Di Giammartino, D. C., Li, J., Kim, D., Kita, K., … Apostolou, E. (2017). Widespread Mitotic Bookmarking by Histone Marks and Transcription Factors in Pluripotent Stem Cells. Cell Reports, 19(7), 1283–1293. https://doi.org/10.1016/J.CELREP.2017.04.067

Martin, C.-A., Murray, J. E., Carroll, P., Leitch, A., Mackenzie, K. J., Halachev, M., … Jackson, A. P. (2016). Mutations in genes encoding condensin complex proteins cause microcephaly through decatenation failure at mitosis. Genes & Development, 30(19), 2158–2172. https://doi.org/10.1101/gad.286351.116

Martinez-Balbbs, M. A., Dey, A., Rabindran, S. K., Ozato, K., & Wu, C. Displacement of Sequence-Specific Transcription Factors from Mitotic Chromatin, 83 Cell § (1995). Retrieved from https://www.cell.com/cell/pdf/0092-8674(95)90231-7.pdf

McGranahan, N., Burrell, R. A., Endesfelder, D., Novelli, M. R., & Swanton, C. (2012, June 1). Cancer chromosomal instability: Therapeutic and diagnostic challenges. EMBO Reports. EMBO Press. https://doi.org/10.1038/embor.2012.61

McKinney, W. (2010). Data Structures for Statistical Computing in Python. In Proceedings of the 9th Python in Science Conference (pp. 56–61). https://doi.org/10.25080/Majora-92bf1922-00a

Molina, O., Vinyoles, M., Granada, I., Roca-Ho, H., Gutierrez-Agüera, F., Valledor, L., … Menendez, P. (2020). Impaired Condensin Complex and Aurora B kinase underlie mitotic and chromosomal defects in hyperdiploid B-cell ALL. Blood. https://doi.org/10.1182/blood.2019002538

Natsume, T., & Kanemaki, M. T. (2017). Conditional Degrons for Controlling Protein Expression at the Protein Level. Annual Review of Genetics, 51(1), 83–102. https://doi.org/10.1146/annurev-genet-120116-024656

Nishimura, K., Fukagawa, T., Takisawa, H., Kakimoto, T., & Kanemaki, M. (2009). An auxin-based degron system for the rapid depletion of proteins in nonplant cells. Nature Methods, 6(12), 917–922. https://doi.org/10.1038/nmeth.1401

Oliphant, T. E. (2006). Guide to NumPy. Retrieved from http://www.trelgol.com

Ono, T., Yamashita, D., & Hirano, T. (2013). Condensin II initiates sister chromatid resolution during S phase. The Journal of Cell Biology, 200(4), 429–441. https://doi.org/10.1083/jcb.201208008

Oser, M. G., Fonseca, R., Chakraborty, A. A., Brough, R., Spektor, A., Jennings, R. B., … Kaelin, W. G. (2019). Cells Lacking the RB1 Tumor Suppressor Gene Are Hyperdependent on Aurora B Kinase for Survival. Cancer Discovery, 9(2), 230–247. https://doi.org/10.1158/2159-8290.CD-18-0389

Perez, F., & Granger, B. E. (2007). IPython: A System for Interactive Scientific Computing. Computing in Science & Engineering, 9(3), 21–29. https://doi.org/10.1109/MCSE.2007.53

Potapova, T., & Gorbsky, G. J. (2017). The Consequences of Chromosome Segregation Errors in Mitosis and Meiosis. Biology, 6(4), 12. https://doi.org/10.3390/biology6010012

Quénet, D., & Dalal, Y. (2014). A long non-coding RNA is required for targeting centromeric protein A to the human centromere. ELife, 3, e03254. https://doi.org/10.7554/eLife.03254

Raccaud, M., Friman, E. T., Alber, A. B., Agarwal, H., Deluz, C., Kuhn, T., … Suter, D. M. (2019). Mitotic chromosome binding predicts transcription factor properties in interphase. Nature Communications, 10(1), 487. https://doi.org/10.1038/s41467-019-08417-5

Ricke, R. M., Jeganathan, K. B., & van Deursen, J. M. (2011). Bub1 overexpression induces aneuploidy and tumor formation through Aurora B kinase hyperactivation. The Journal of Cell Biology, 193(6), 1049–1064. https://doi.org/10.1083/jcb.201012035

Rohrberg, J., Van de Mark, D., Amouzgar, M., Lee, J. V., Taileb, M., Corella, A., … Goga, A. (2020). MYC Dysregulates Mitosis, Revealing Cancer Vulnerabilities. Cell Reports, 30(10), 3368–3382.e7. https://doi.org/10.1016/J.CELREP.2020.02.041

Ruppert, J. G., Samejima, K., Platani, M., Molina, O., Kimura, H., Jeyaprakash, A. A., … Earnshaw, W. C. (2018). HP1a targets the chromosomal passenger complex for activation at heterochromatin before mitotic entry. The EMBO Journal, 37(6), 97677. https://doi.org/10.15252/embj.201797677

Salimian, K. J., Ballister, E. R., Smoak, E. M., Wood, S., Panchenko, T., Lampson, M. A., & Black, B. E. (2011). Feedback Control in Sensing Chromosome Biorientation by the Aurora B Kinase. Current Biology, 21, 1158–1165. https://doi.org/10.1016/j.cub.2011.06.015

Samejima, K., Booth, D. G., Ogawa, H., Paulson, J. R., Xie, L., Watson, C. A., … Earnshaw, W. C. (2018). Functional analysis after rapid degradation of condensins and 3D-EM reveals chromatin volume is uncoupled from chromosome architecture in mitosis. Journal of Cell Science, 131(4). https://doi.org/10.1242/jcs.210187

Samoshkin, A., Arnaoutov, A., Jansen, L. E. T., Ouspenski, I., Dye, L., Karpova, T., … Strunnikov, A. (2009). Human Condensin Function Is Essential for Centromeric Chromatin Assembly and Proper Sister Kinetochore Orientation. PLoS ONE, 4(8), e6831. https://doi.org/10.1371/journal.pone.0006831

Sanjana, N. E., Shalem, O., & Zhang, F. (2014). Improved vectors and genome-wide libraries for CRISPR screening. Nature Methods, 11(8), 783–784. https://doi.org/10.1038/nmeth.3047

Schober, P., & Schwarte, L. A. (2018). Correlation coefficients: Appropriate use and interpretation. Anesthesia and Analgesia, 126(5), 1763–1768. https://doi.org/10.1213/ANE.0000000000002864

Shimomura, H., Sasahira, T., Nakashima, C., Kurihara-Shimomura, M., & Kirita, T. (2019). Non-SMC Condensin I Complex Subunit H (NCAPH) Is Associated with Lymphangiogenesis and Drug Resistance in Oral Squamous Cell Carcinoma. Journal of Clinical Medicine, 9(1), 72. https://doi.org/10.3390/jcm9010072

Sotillo, R., Hernando, E., Díaz-Rodríguez, E., Teruya-Feldstein, J., Cordón-Cardo, C., Lowe, S. W., & Benezra, R. (2007). Mad2 overexpression promotes aneuploidy and tumorigenesis in mice. Cancer Cell, 11(1), 9. https://doi.org/10.1016/J.CCR.2006.10.019

Sutani, T., Sakata, T., Nakato, R., Masuda, K., Ishibashi, M., Yamashita, D., … Shirahige, K. (2015). Condensin targets and reduces unwound DNA structures associated with transcription in mitotic chromosome condensation. Nature Communications, 6(1), 7815. https://doi.org/10.1038/ncomms8815

Takemoto, A., Murayama, A., Katano, M., Urano, T., Furukawa, K., Yokoyama, S., … Kimura, K. (2007). Analysis of the role of Aurora B on the chromosomal targeting of condensin I. Nucleic Acids Research, 35(7), 2403–2412. https://doi.org/10.1093/nar/gkm157

Tang, A., Gao, K., Chu, L., Zhang, R., Yang, J., & Zheng, J. (2017). Aurora kinases: novel therapy targets in cancers. Oncotarget (Vol. 8). Retrieved from www.impactjournals.com/oncotarget/

Tate, J. G., Bamford, S., Jubb, H. C., Sondka, Z., Beare, D. M., Bindal, N., … Forbes, S. A. (2019). COSMIC: the Catalogue Of Somatic Mutations In Cancer. Nucleic Acids Research, 47(D1), D941–D947. https://doi.org/10.1093/nar/gky1015

Taylor, A. M., Shih, J., Ha, G., Cherniack, A. D., Beroukhim, R., & Meyerson Correspondence, M. (2018). Genomic and Functional Approaches to Understanding Cancer Aneuploidy. https://doi.org/10.1016/j.ccell.2018.03.007

Teves, S. S., An, L., Hansen, A. S., Xie, L., Darzacq, X., & Tjian, R. (2016). A dynamic mode of mitotic bookmarking by transcription factors. ELife, 5(NOVEMBER2016), 1–24. https://doi.org/10.7554/eLife.22280

Torabi, B., Flashner, S., Beishline, K., Sowash, A., Donovan, K., Bassett, G., & Azizkhan-Clifford, J. (2018). Caspase cleavage of transcription factor Sp1 enhances apoptosis. Apoptosis, 23(1), 65–78. https://doi.org/10.1007/s10495-017-1437-4

van der Lelij, P., Lieb, S., Jude, J., Wutz, G., Santos, C. P., Falkenberg, K., … Petronczki, M. (2017). Synthetic lethality between the cohesin subunits STAG1 and STAG2 in diverse cancer contexts. ELife, 6. https://doi.org/10.7554/eLife.26980

van der Walt, S., Colbert, S. C., & Varoquaux, G. (2011). The NumPy Array: A Structure for Efficient Numerical Computation. Computing in Science & Engineering, 13(2), 22–30. https://doi.org/10.1109/MCSE.2011.37

Venter, J. C., Adams, M. D., Myers, E. W., Li, P. W., Mural, R. J., Sutton, G. G., … Zhu, X. (2001). The sequence of the human genome. Science (New York, N.Y.), 291(5507), 1304–1351. https://doi.org/10.1126/science.1058040

Virtanen, P., Gommers, R., Oliphant, T. E., Haberland, M., Reddy, T., Cournapeau, D., … van Mulbregt, P. (2020). SciPy 1.0: fundamental algorithms for scientific computing in Python. Nature Methods, 17(3), 261–272. https://doi.org/10.1038/s41592-019-0686-2

Walther, N., Julius Hossain, M., Politi, A. Z., Koch, B., Kueblbeck, M., Ødegård-Fougner, Ø., … Ellenberg, J. (2018). A quantitative map of human Condensins provides new insights into mitotic chromosome architecture. J. Cell Biol, 217, 2309. https://doi.org/10.1083/jcb.201801048

Wang, H.-Z., Yang, S.-H., Li, G.-Y., & Cao, X. (2018). Subunits of human condensins are potential therapeutic targets for cancers. Cell Division, 13(1), 2. https://doi.org/10.1186/s13008-018-0035-3

Weaver, B. A., & Cleveland, D. W. (2006). Does aneuploidy cause cancer? Current Opinion in Cell Biology, 18(6), 658–667. https://doi.org/10.1016/j.ceb.2006.10.002

Weiler, S. M. E., Pinna, F., Wolf, T., Lutz, T., Geldiyev, A., Sticht, C., … Breuhahn, K. (2017). Induction of Chromosome Instability by Activation of Yes-Associated Protein and Forkhead Box M1 in Liver Cancer. Gastroenterology, 152(8), 2037–2051.e22. https://doi.org/10.1053/J.GASTRO.2017.02.018

Wilkinson, R. W., Odedra, R., Heaton, S. P., Wedge, S. R., Keen, N. J., Crafter, C., … Green, S. (2007). AZD1152, a Selective Inhibitor of Aurora B Kinase, Inhibits Human Tumor Xenograft Growth by Inducing Apoptosis. Clinical Cancer Research, 13(12), 3682–3688. https://doi.org/10.1158/1078-0432.CCR-06-2979

Woodward, J., Taylor, G. C., Soares, D. C., Boyle, S., Sie, D., Read, D., … Wood, A. J. (2016). Condensin II mutation causes T-cell lymphoma through tissue-specific genome instability. Genes and Development, 30(19), 2173–2186. https://doi.org/10.1101/gad.284562.116

Zhang, Q., Su, R., Shan, C., Gao, C., & Wu, P. (2018). Non-SMC Condensin I Complex, Subunit G (NCAPG) is a Novel Mitotic Gene Required for Hepatocellular Cancer Cell Proliferation and Migration. Oncology Research Featuring Preclinical and Clinical Cancer Therapeutics, 26(2), 269–276. https://doi.org/10.3727/096504017X15075967560980

Zhang, Y., Liu, F., Zhang, C., Ren, M., Kuang, M., Xiao, T., … Cheng, S. (2020). Non-SMC Condensin I Complex Subunit D2 Is a Prognostic Factor in Triple-Negative Breast Cancer for the Ability to Promote Cell Cycle and Enhance Invasion. The American Journal of Pathology, 190(1), 37–47. https://doi.org/10.1016/J.AJPATH.2019.09.014

