## Supplemental Table - list of primer sequences for "Transcription factor Sp1 regulates mitotic fidelity through Aurora B kinase-mediated condensin I localization"

| Primer Name | Seq (5' - 3') |
|---|---|
| sgSp1 F | CACCGCATGGATGAAATGACAGCTG |
| sgSp1 R | AAACCAGCTGTCATTTCATCCATGC |
| H2B-mCherry F | TATTACCATGGTGATGCGGTTTTGG |
| H2B-mCherry R | GAGGTTGATTTTACTTGTACAGCTCGT |
| pLENTI (for mCherry H2B) F | GTACAAGTAAAATCAACCTCTGGATTACA |
| pLENTI (for mcherry H2B) R | ACCGCATCACCATGGTAATAGCGATGAC |
| pLZS Sp1 F | GCCCGAGGGATCCGCTGCTGCC |
| pLZS Sp1 R | CTTTAGAGTCCATGGTGGCAGCTG |
| pCDNA eGFP-mAID F | TGCCACCATGGACTCTAAAGAGAAGTCAG |
| pcDNA eGFP-mAID R | CAGCAGCGGATCCCTCGGGCCC |

Supplementary Table 1: List of Primers used in this study
